# ClsDiff-AMP30: Generating Antimicrobial Peptides by a Classifier Guidance Noise Predictor

**DOI:** 10.1101/2025.11.25.690620

**Authors:** Jielu Yan, Jianxiu Cai, Yifan Li, Zhongyao Lin, Weizhi Xian, Xuekai Wei, Iun Fan Lei, Mingliang Zhou, François-Xavier Campbell-Valois, Shirley W. I. Siu

**Author notes:** These authors contributed equally to this work (Jielu Yan, Jianxiu Cai).

## Abstract

Antimicrobial peptides (AMPs) represent a promising therapeutic strategy to combat the increasing challenge of multidrug-resistant pathogens, a crisis intensified by the overuse of conventional antibiotics. In addition to their broad-spectrum antimicrobial activity, low toxicity, and reduced propensity for resistance development, AMPs offer significant advantages over traditional antibiotic therapies. However, the discovery of novel AMPs through biological experiments remains constrained by high costs, labor-intensive workflows, and time-consuming procedures, underscoring the urgent need for *in silico* computational methods to design AMP sequences. Notably, shorter AMPs (*≤* 30 residues) demonstrate superior antimicrobial efficacy, improved structural stability, and minimal cytotoxicity toward human cells. To address these challenges, we present a classifier-guided diffusion framework specialized for generating AMPs shorter than 30 residues (ClsDiff-AMP30). The architecture integrates two interdependent submodels, including a noisy AMP classifier that evaluates AMP likelihood at intermediate denoising steps and a noise predictor guided by classifier-derived probability scores, dynamically adjusted via a self-optimized coefficient to modulate guidance strength. ClsDiff-AMP30 achieves a validation accuracy of 66% across 10,000 synthesized sequences by a self-developed AMP classifier. Furthermore, wet lab experiments demonstrated that all 11 selected sequences exhibited high antimicrobial activity against at least one of the three tested bacterial strains and low hemolytic activity.

## 1 Introduction

With the growing threat of multidrug resistance in pathogens caused by the overuse of conventional antibiotics, the design of novel antibiotics has emerged as a critical research focus. Antimicrobial peptides (AMPs) exhibit broad-spectrum antimicrobial activities, including antibacterial, antifungal, antiviral, and anticancer activities [1], while demonstrating lower toxicity and fewer side effects [2], rapid action [3], and a reduced propensity to induce resistance [4]. These advantages position AMPs as promising candidates to replace traditional antibiotics in addressing multidrug resistance. However, conventional wet-lab experiments face limitations such as being labor intensive, time consuming, and resource inefficient [5]. Additional challenges include low antimicrobial efficacy, poor stability, and high hemolytic activity, which hinder clinical translation [6]. Moreover, traditional methods struggle to screen and analyse the vast combinatorial space of potential peptide sequences in the era of big data [7]. Consequently, there is an urgent need to develop *in silico* tools that leverage computational efficiency to accelerate AMP discovery. Such tools can systematically analyse physicochemical parameters influencing antibacterial activity, protease stability, cytotoxicity, and hemolytic activity, thereby elucidating molecular mechanisms [7, 8].

Numerous studies have explored AMP generation via diverse deep learning architectures, including recurrent neural networks (RNNs), long short-term memory networks (LSTMs), generative adversarial networks (GANs), variational autoencoders (VAEs), adversarial autoencoders (AAEs), and transformer networks [9–12]. Müller *et al.* [13] proposed a generative RNN framework for combinatorial *de novo* peptide design, which uses helical AMPs with learned contextual features. The model achieved 82% prediction accuracy for active AMPs (versus 65% for random sequences with matched amino acid distributions) while generating sequences closer to natural amphipathic helices than manual designs. Wang *et al.* [14] developed a bidirectional LSTM classifier paired with an LSTM-based generator to design short AMPs targeting *E. coli*, demonstrating enhanced antibacterial activity prediction. Ferrell *et al.* [15] introduced AMP-GAN, a GAN trained on 500,000 non-AMP and 8,000 AMP sequences. Through molecular dynamics screening of the generated sequences, six candidates were synthesized, three of which exhibited validated bacterial growth inhibition. To achieve this success, Van Oort *et al.* [16] subsequently developed AMPGAN v2, a bidirectional conditional GAN (BiCGAN) that incorporates conditioning variables to control generation while maintaining AMP characteristics across diverse outputs. Dean *et al.* [17] proposed PepVAE, a variational autoencoder that integrates sequence data and minimum inhibitory concentrations (MICs) to generate novel AMPs with experimentally validated activity. Renaud *et al.* [18] systematically compared five architectures (RNN, RNN with attention, AAE, transformer) across three latent space dimensions (32, 64, 128). Their analysis of reconstruction accuracy, generation capability, and interpretability revealed that most models partition latent spaces into regions correlated with AMP likelihood and physicochemical properties. This enables targeted sampling of AMPs with optimized characteristics through latent space manipulation.

However, most on-the-shelf approaches have limitations. RNNs struggle with long-term dependencies due to the vanishing gradient problem, which affects their ability to capture long-range interactions in sequence data [19–21]. LSTMs, while improved over RNNs for capturing long-term dependencies, are computationally expensive and can be prone to overfitting with complex sequence data [22, 23]. GANs can suffer from mode collapse and unstable training dynamics, which makes generating high-quality and diverse samples challenging [24, 25]. VAEs may produce blurry outputs because of the discretization of the latent space, and their performance can be limited by the capacity of the decoder network [26, 27]. Compared with recent advancements in diffusion models, AAEs have difficulty modelling complex distributions and generating high-fidelity samples, especially [28, 29]. Transformers, while powerful for sequence-to-sequence tasks, can be less efficient for generative modelling tasks that require fine-grained control over the data distribution [30–32]. In contrast, diffusion models generate a sequence from arbitrary Gaussian noise through iterative refinement. These models operate under the assumption that the feature vector of a given sequence can be systematically perturbed by additive noise (i.e., Gaussian noise) at each timestep. After a total of *T* timesteps, the original input feature vector becomes fully corrupted into Gaussian noise. Conversely, the generation process involves using a trained model to iteratively reconstruct the denoised input at timestep *t* −−1 from its noisy counterpart at timestep *t*, starting from random Gaussian noise. In summary, diffusion models provide a flexible framework capable of synthesizing high-quality samples across diverse data modalities. Inspired by thermodynamic diffusion principles, they learn to reverse the noise-adding process, thereby enhancing sample diversity and fidelity [33]. Unlike GANs, which rely on discrete adversarial training steps, noise predictors employ a continuous generative process that improves stability and controllability. Furthermore, their compatibility with diverse likelihood functions enables greater flexibility in modelling complex data distributions.

In this study, we proposed ClsDiff-AMP30, a classifier-guided diffusion model trained on AMPs with fewer than 30 residues. ClsDiff-AMP30 comprises two key components: a noise predictor that generates novel candidate AMPs and a noisy classifier that guides the noise predictor to produce higher-scoring AMP candidates. During training, the noise predictor and noise classifier were trained separately, both of which used noisy tokens as input features. The noise predictor predicts noise on the AMP dataset and then computes the corresponding noise tokens for the previous timestep through a series of formulas. The noisy classifier predicts an AMP probability score ranging from 0 to 1 at each timestep on both the AMP and non-AMP datasets. In the generation phase, ClsDiff-AMP30 synthesizes sequences directly from a Gaussian distribution, leveraging the AMP probability scores predicted by the noisy classifier to iteratively refine the noisy tokens produced by the noise predictor. To evaluate the performance of ClsDiff-AMP30, we trained a random forest (RF) classifier to distinguish AMPs from non-AMPs. When generating 10,000 peptides, the model achieved an accuracy of 66%. Additionally, we conducted a series of experiments to evaluate the quality of the generated AMPs by comparing them with real AMPs, randomly generated sequences, and random distinct peptides. To assess the robustness of ClsDiff-AMP30, we varied the input noise distribution; tested Gaussian, uniform, and gamma noise; explored the impact of training dataset size on model performance; and evaluated whether the noisy classifier enhanced the model’s generative capability. We subsequently selected 12 out of the 130 candidate sequences, analysed their 3D structures and identified three motifs for further validation. Finally, we conducted wet lab experiments to evaluate the 11 shortlisted sequences against three bacterial strains. The longest sequence from the original set of 12 was excluded to reduce costs. All 11 tested sequences demonstrated high antimicrobial activity and low hemolytic activity.

## 2 Materials and Methods

### 2.1 Data collection

We curated three specialized datasets to develop and validate our generative framework. The first dataset is the AMP dataset for generative model development. The second dataset is the non-AMP dataset required for training the auxiliary classifier that guides and scores the generated sequences. The third dataset is the PeptideAtlas dataset, an independent diverse peptide set used for comparative analysis of model-generated AMPs.

### AMP dataset

The AMP sequences were downloaded from four published online databases: DBAASP [34], DRAMP [35], APD3 [36], and DBAMP [37]. In total, 39,684 AMPs were downloaded, and a preprocessing procedure was performed for generation purposes. We removed all sequences that contained symbols other than the 20 natural amino acids, had lengths outside the range of 5–30 residues and had duplicate sequences. Finally, we obtain 27,180 AMPs as our final AMP dataset.

### Non-AMP dataset

Since UniProt [38] does not contain sufficient non-AMP sequences with the same length distribution as the AMP sequences do, we constructed the non-AMP dataset by computationally generating synthetic negative sequences via the method proposed by Yan *et al.* [39]. This non-AMP dataset is size-matched to the positive AMP dataset to ensure class balance and is aligned with its length distribution, thereby mitigating potential biases arising from divergent sequence–length profiles.

### PeptideAtlas dataset

We collected all the distinct peptides from the Peptide Atlas database [40] and obtained 3,979,055 peptides in total. After rigorous preprocessing (mirroring the AMP pipeline) and cross-dataset duplicate removal, the final set consisted of 3,706,665 nonredundant sequences.

Table 1 summarizes the composition of the AMP, non-AMP, and peptide datasets, including their sources (database names or generation methods), original sizes, and postprocessed counts. Further-more, we analysed the AMP and non-AMP datasets, as well as the naturally occurring peptides from PeptideAtlas, focusing on the sequence length distributions and amino acid compositions. As illustrated in Figure 1a, the AMP and non-AMP datasets exhibit identical length distributions, whereas the peptide dataset deviates significantly, with over 30% of its sequences falling within the 11–15 residue range. Regarding the amino acid composition shown in Figure 1b, the non-AMP dataset shows a uniform distribution across all 20 natural amino acids. In contrast, the AMP dataset is enriched in residues *R* (arginine), *L* (leucine), and *K* (lysine) but depleted in *M* (methionine), *E* (glutamic acid), and *D* (aspartic acid). The peptide dataset demonstrates distinct biases, elevated proportions of *L* (leucine), *E* (glutamic acid), and *S* (serine), alongside reduced frequencies of *W* (tryptophan), *C* (cysteine), and *M* (methionine).

**Figure 1:**
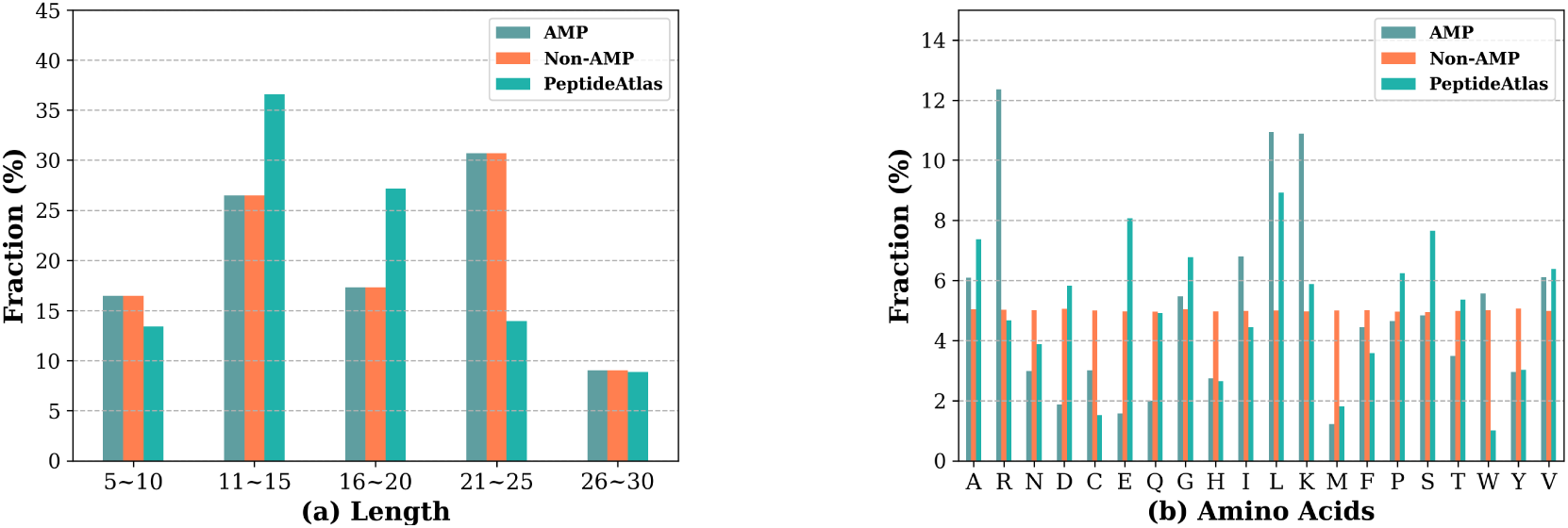
Distribution of sequence length (a) and amino acid composition (b) of the AMP, non-AMP, and PeptideAtlas datasets.

**Table 1:** Summary of the AMP, synthetic non-AMP, and natural peptide datasets.

|  | AMP | Non-AMP | PeptideAtlas |
| --- | --- | --- | --- |
| **Database source or**<br>**Generation Method** | DBAASP v3 [34] | Random generation [39] | PeptideAtlas [40] |
|  | DRAMP [35] |  |  |
|  | APD3 [36] |  |  |
|  | DBAMP [37] |  |  |
| **Original number** | 39,684 | 27,180 | 3,979,590 |
| **After preprocessing** | 27,180 | – | 3,706,665 |

In addition, we analysed the AMP, non-AMP, and peptide datasets, including the length and amino acid compositions. As shown in Figure 1, the length distributions of the AMP dataset are the same as those of the non-AMP dataset. However, the length distribution of our additional dataset, i.e., the peptide dataset, is different from those of the AMP and non-AMP datasets. More than 30% of peptide lengths are in the range of 11–15. For amino acid composition, the non-AMP dataset contains the same composition for all 20 different natural amino acids. The AMP dataset has more “R”, “L”, and “K” values and contains fewer “M”, “E”, and “D” values. The peptide dataset contains more amino acids that are short for “L”, “E”, and “S” and has fewer amino acids of “W”, “C”, and “M”.

### 2.2 Methods

#### 2.2.1 Problem formulation

For the development of the AMP generative model, the problem can be formulated as follows: Given an arbitrary Gaussian noise vector, a trained generative diffusion model generates the corresponding

AMP sequence. A peptide sequence of length *N* is represented as a series of *N* characters (*S_i_* ∈ {1, 2, 3*, …, N* }), where each character corresponds to a natural amino acid:

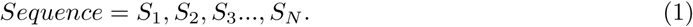

Prior to deep learning, an input sequence is first converted into a series of tokens via a token encoding method. Here, the sequence token dictionary shown in Eq. (2) is employed to map the 20 natural amino acids to integer values ranging from 1–20. To standardize sequence lengths, the integer value 24 is used as a padding token for sequences shorter than the maximum length, whereas the end-of-sequence is denoted by the end token 25.

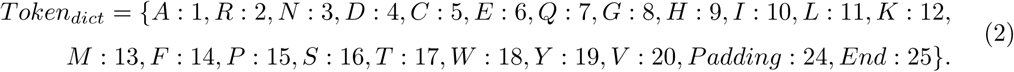

As shown in Eq. (3), a token vector *Token* is an input feature vector whose size is the maximum length of the training AMPs plus one. Each token *F_i_* in *Token*, where *i* ∈ {1, 2, 3*, …, M* }, denotes the token value of the corresponding amino acid in the i-th position of the input sequence converted via *Token_dict_*. To standardize all the input features to the range of [-1, 1], we applied a normalization operation to each sequence by first dividing the values by 12.5 and then subtracting 1.

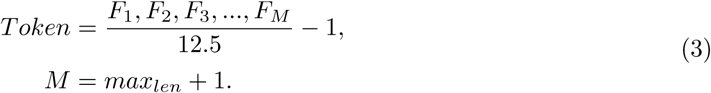

The proposed generative diffusion framework begins with the *noise-adding procedure*. Assuming that the clean input token *Token*_0_ can be perturbed by Gaussian noise of varying scales in a step-wise manner, the noisy token *Token_T_* becomes indistinguishable from pure Gaussian noise after a sufficiently large number of timesteps *T* . Mathematically, each intermediate noise token *Token_t_* (for *t* ∈ {1, 2, 3*, …, T* }) is computed via a predefined formula, forming a Markov chain that progressively obfuscates the original signal.

The inversion of this diffusion process is enabled by the noise predictor, a trained deep learning model that predicts the added noise at each timestep *t* for a given *Token_t_*, and outputs the corresponding *Token_t−_*_1_. Therefore, starting from an arbitrarily sampled Gaussian noise *Token_T_* and iteratively denoising from *t* = *T* to *t* = 1, the model generates the clean token *Token*_0_. The process of training a noise predictor model to predict the added noise at an arbitrary timestep *t* from *Token_t_* is referred to as the *training procedure of the noise predictor*.

To increase the efficiency of the generative model in producing high-quality AMP sequences, an extra noisy classifier model is employed. This classifier uses as input the timestep *t* and the *t*-th step noised token *Token_t_*, outputting a probability score *ŷ* that quantifies the likelihood of the sequence being a valid AMP or not.

Furthermore, from the noisy classifier model, the guidance information *Token^guidance^* between the guidance label and the input *Token_t_* can be extracted, which is used to guide the generated *Token_t_* toward sequences with higher predicted AMP scores at every timestep. Following denoising, the final generated *Token*_0_ is mapped to an amino acid sequence via the token dictionary, completing the sequence generation procedure.

In summary, the four procedures–the noise-adding procedure, the training procedure of the noise predictor, the training procedure of the noisy classifier, and the sequence generation procedure–constitute the core of the proposed generative diffusion framework and are discussed in detail.

#### 2.2.2 Proposed generative diffusion framework

##### Noise-adding procedure

The noise-adding procedure, also known as the forward diffusion process, is mathematically defined by a closed-form formula, as shown in Eq. (4), which enables the efficient computation of a corrupted token *Token_t_* at any timestep *t*.

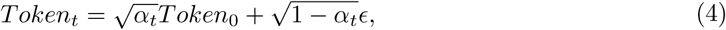

where *Token*_0_ represents the original token. *Token_t_* is the noised token derived from *Token*_0_ by applying *t* increments of standard Gaussian noise *ɛ* as defined in Eq. (5), which is sampled anew at each timestep.

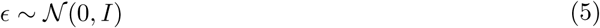

The cumulative noise decay coefficient, denoted as *α_t_* and defined in Eq. (6), exhibits monotonic decay with increasing timestep *t*.

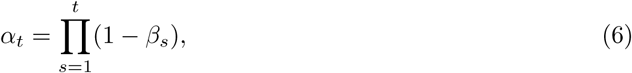

where *β* is the noise schedule that controls the rate of diffusion. Here, *β_t_* at timestep *t* is parameterized such that its minimum and maximum values are given by *β_min_* and *β_max_*, respectively.

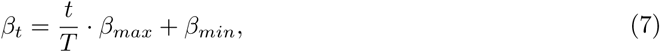

where *T* denotes the total number of timesteps. In this work, *β_min_* and *β_max_* are set to 0.0001 and 0.02, respectively.

#### Training procedure of the noise predictor

The noise predictor, a specialized deep learning architecture, is engineered to iteratively estimate the denoised token *Token_t−_*_1_ from its corrupted counterpart *Token_t_*. Through this reverse diffusion process, the trained model can gradually reconstruct realistic tokens starting from arbitrary Gaussian noise samples. Specifically, given any initial Gaussian noise input and a trained noise predictor, we can generate synthetic tokens resembling the original training data through stepwise denoising operations.

The noise prediction process can be formalized as in Eq. (8):

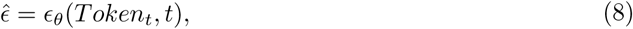

where *ɛ_θ_* denotes the noise predictor model parameterized by weights *θ*. During training, the timestep *t* is sampled uniformly from the set as defined in Eq. (9):

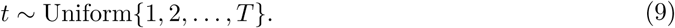

The loss function of the noise predictor L*_diff_* is shown in Eq. (10), where the noise predictor learns to denoise *Token_t_* by minimizing L*_diff_* .

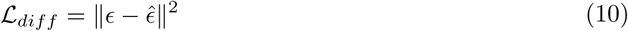

The complete training procedure of the noise predictor is displayed in Algorithm 1.

##### Algorithm 1 Noise Predictor Training Procedure

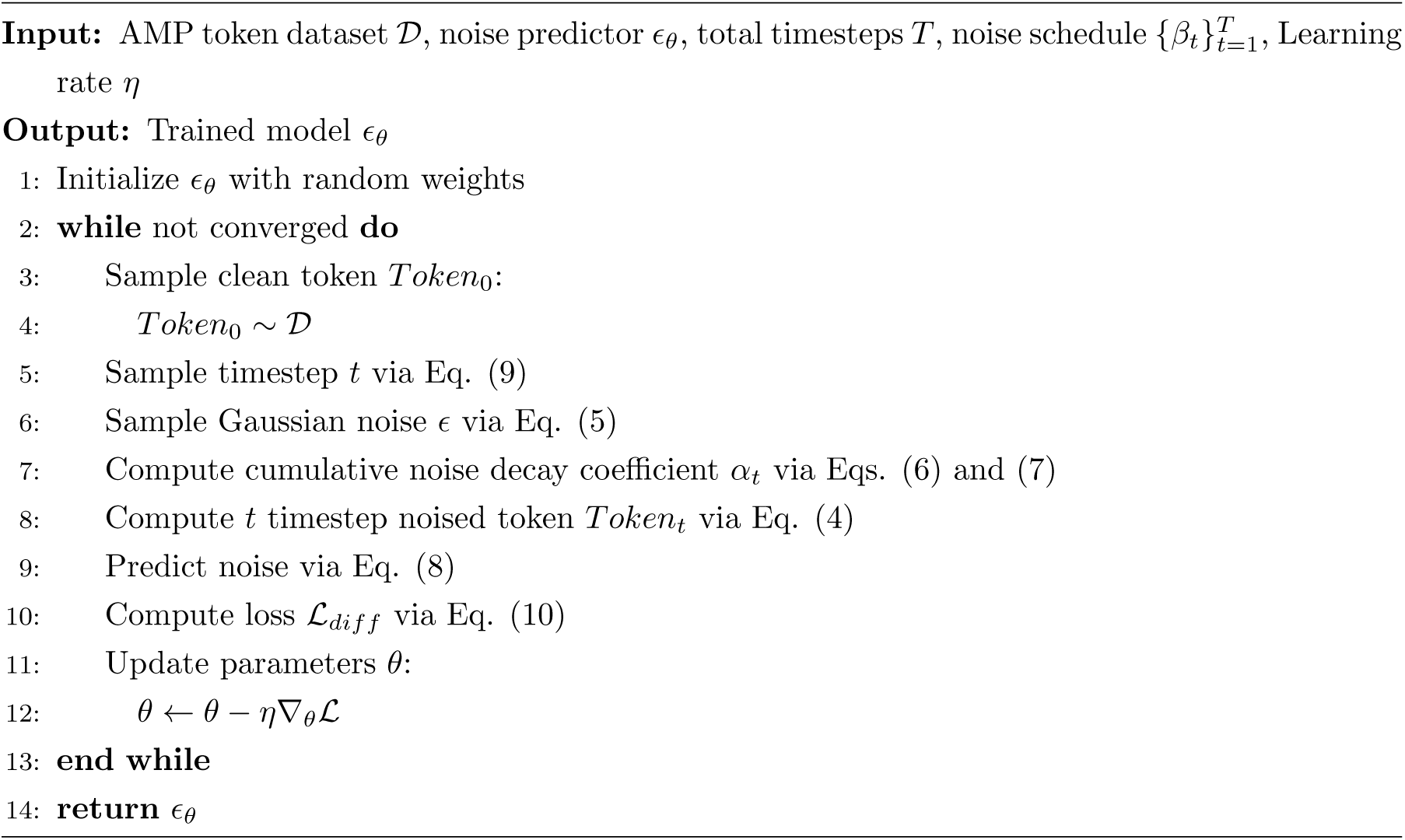

#### Training procedure of the noisy classifier

To improve the generation power of the noise predictor, a noisy classifier model is incorporated to guide the generation of sequences with higher AMP probabilities. The noisy classifier model predicts two classes: AMP (label 1) and non-AMP (label 0). The classification function is defined as:

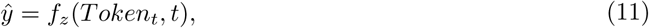

where *f_z_* represents the noisy classifier model, where *z* denotes the set of all the weights of the noisy classifier model and where *ŷ* denotes the AMP probability score generated by *f_z_*. *Token_t_* and *t* are taken as inputs to the noisy classifier model, and *ŷ* is computed. In addition, the loss function used to train the noisy classifier model is defined in Eq. (12):

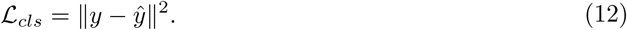

#### Sequence generation procedure

Sequence generation is accomplished through the reverse diffusion process starting from a maximally perturbed token *Token_T_* . The generation process iteratively refines *ClsToken_t−_*_1_ from *Token_t_* by jointly using the trained noise predictor *ɛ_θ_* and the trained noisy classifier *f_z_* models, where *ClsToken_t−_*_1_ represents *Token_t−_*_1_ adjusted by the classifier guidance.

Specifically, to obtain the classifier guidance token *ClsToken_t−_*_1_, the *distance* between the given label *y* and the AMP score predicted by the noisy classifier model is computed via Eq. (13):

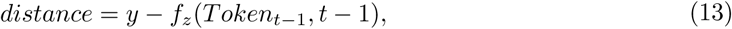

where the guidance label *y* is set to 1 to optimize for an AMP. Furthermore, 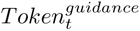 represents the guidance, which is the gradient extracted from the noisy classifier model between the *distance* and *Token_t−_*_1_, as defined in Eq. (14):

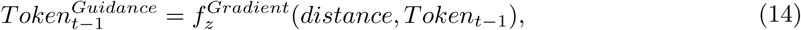

Consequently, the classifier guidance token *ClsToken_t−_*_1_ is computed as the sum of *Token_t−_*_1_ and 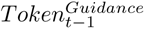 modulated by the guidance scale *λ*:

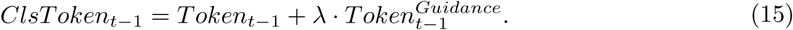

In the last step of reverse diffusion, the unscaled token 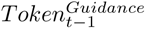 obtained from Eq. (16) is mapped to an amino acid sequence via the token dictionary given in Eq. (2):

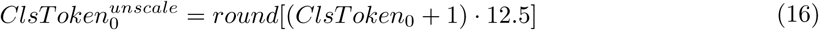

The reverse diffusion step is illustrated in Algorithm 2, and the complete sequence generation procedure is presented in Algorithm 3.

##### Algorithm 2 Generate *Token_t−_*_1_ from *Token_t_*

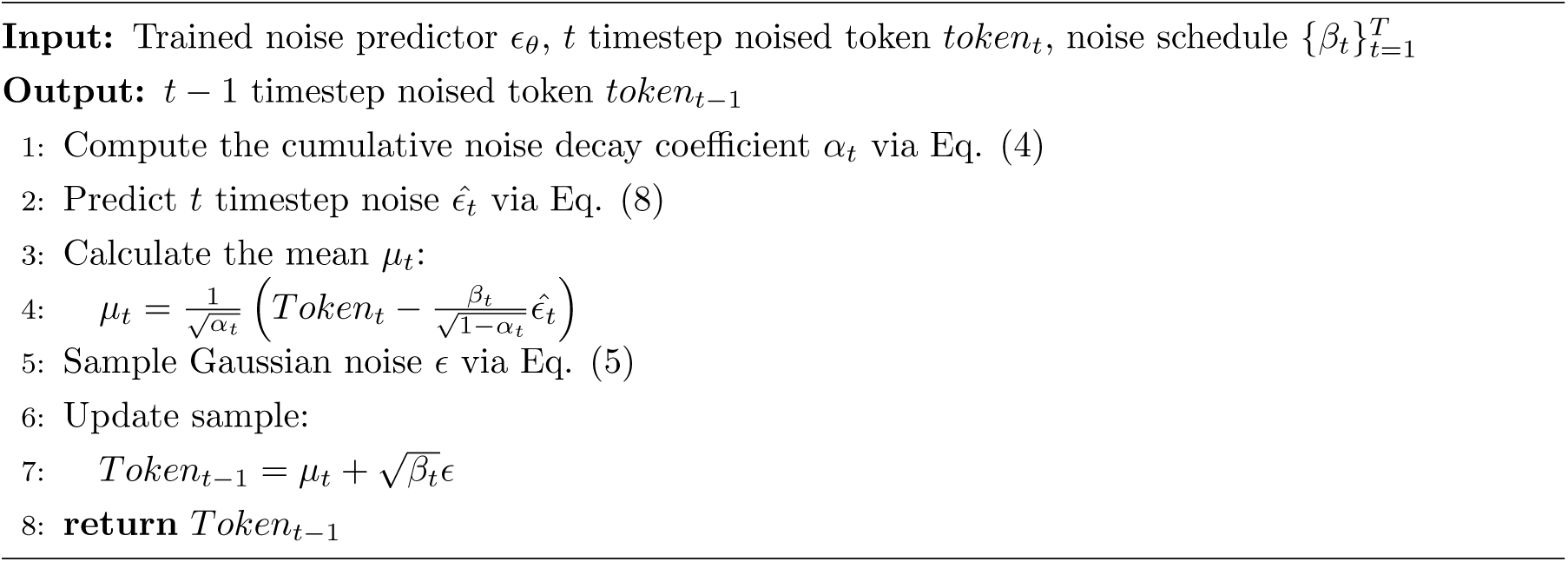

##### Algorithm 3 Sequence generation procedure

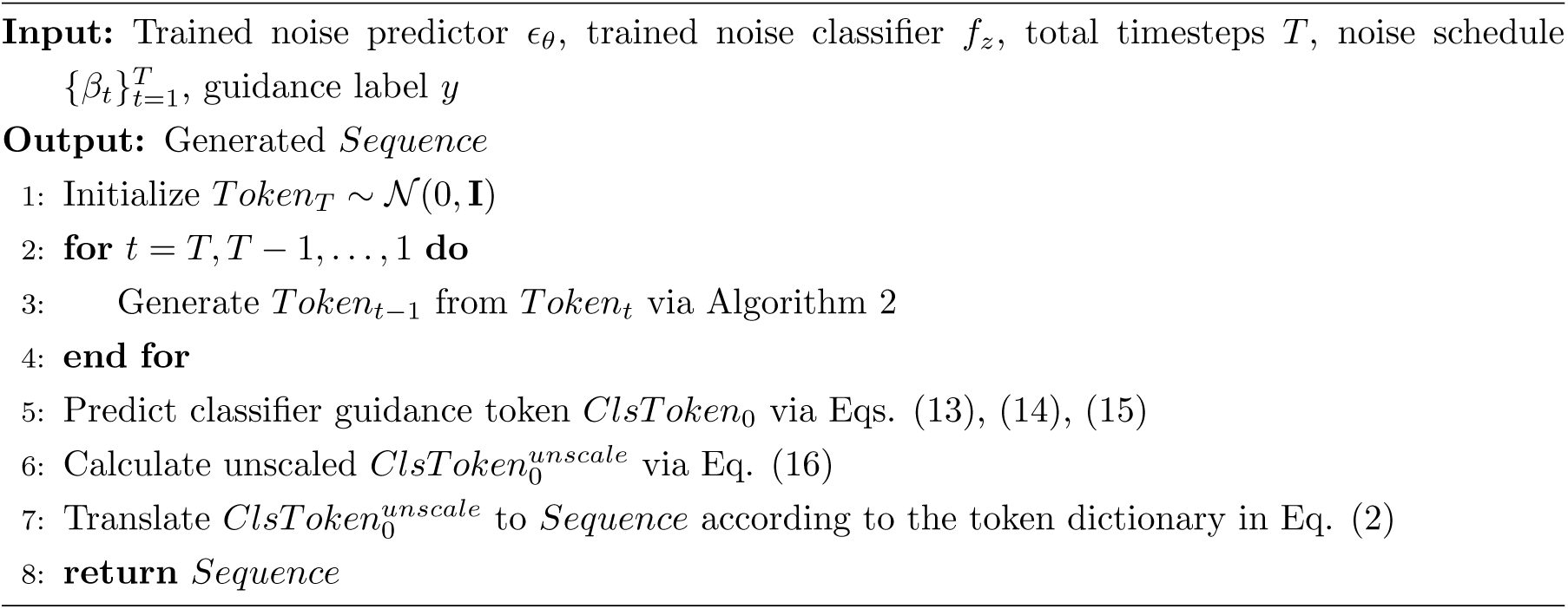

#### 2.2.3 Classifier guidance diffusion model (ClsDiff-AMP30)

To generate short-length AMP sequences, we propose a classifier guidance diffusion model (named ClsDiff-AMP30) for sequence lengths ranging from 5–30. The ClsDiff-AMP30 model consists of two submodels, namely, the noise predictor model to predict the added noise and the noise classifier model to guide the generated tokens with high AMP scores. The input sequences are first converted into clean tokens, and then for each clean token, the noise token is computed for each timestep up to *T* timesteps. The noise predictor generates the previous timestep-noised token from an arbitrary timestep-noised token by deleting the predicted added noise of the corresponding timestep. The novel classifier model guides the generation of tokens to obtain higher AMP scores. In the training procedure, the two submodels utilized *t* timestep noise tokens *Token_t_* as inputs and *t* − 1 timestep noise tokens *Token_t−_*_1_ as outputs.

#### ClsDiff-AMP30

As shown in the upper left part of Figure 2, with *Token_t_* as the input, we compute the *t* − 1 timestep noise token *Token_t−_*_1_ from the *t* timestep noise token *Token_t_* via the denoising model, which is followed by Algorithm 2. The denoised model generates *t* timestep-added noise *ɛ̂_t_* from the *t* timestep-noised token *Token_t_*, and its architecture is shown in Figure 3 and explained in detail in the next part.

**Figure 2:**
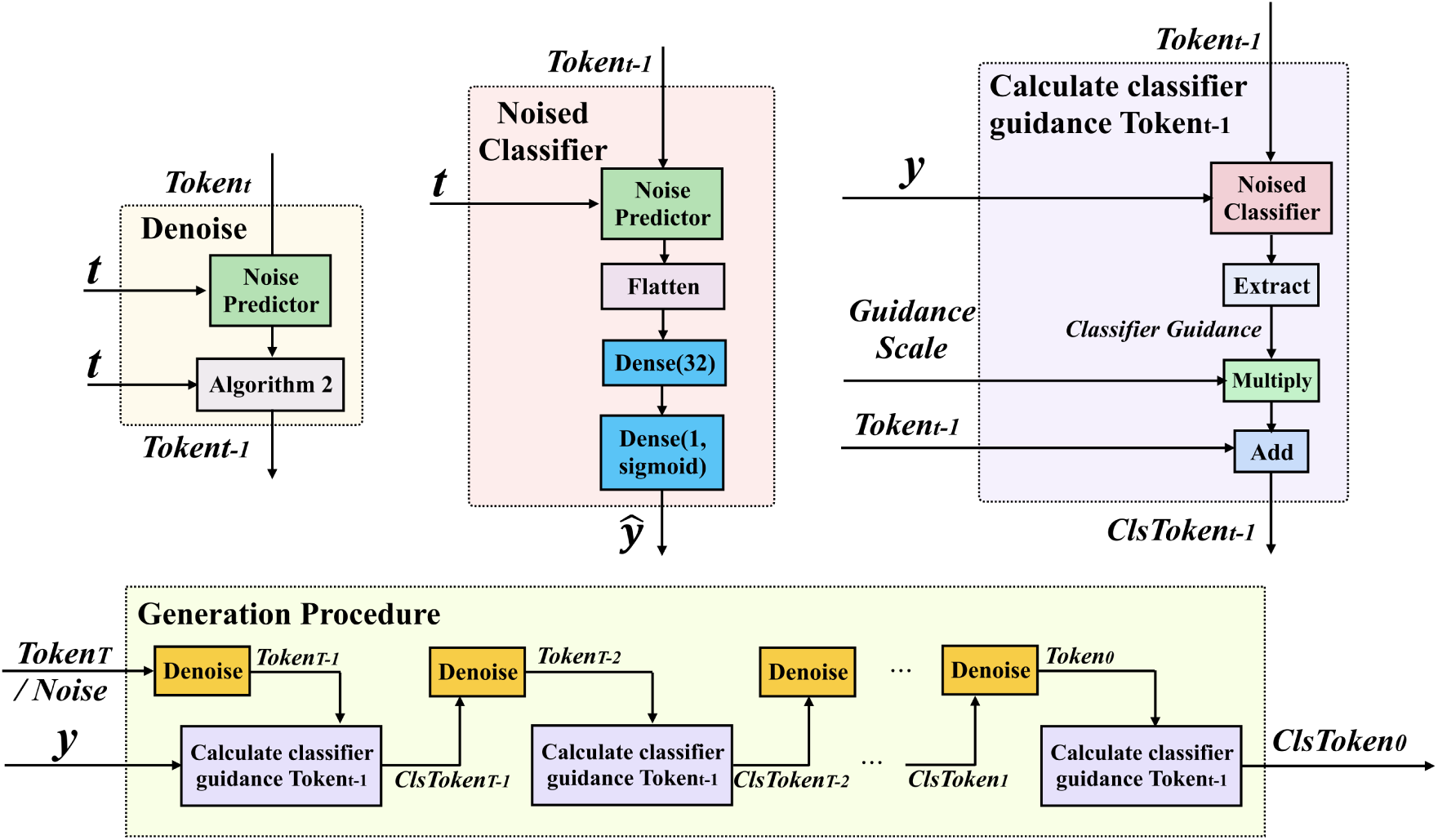
The ClsDiff-AMP30 model. The upper left module is the denoise module, which computes *Token_t−_*_1_ from *Token_t_*. The noisy predictor is shown in Figure 3. The upper middle module illustrates the noise classifier architecture, and the upper right module shows how to calculate the classifier guidance token *ClsToken_t_* from the input *Token_t_*. The bottom module presents the generation procedure of the classifier guidance clean token *ClsToken*_0_ from an input Gaussian noise and a guidance label.

**Figure 3:**
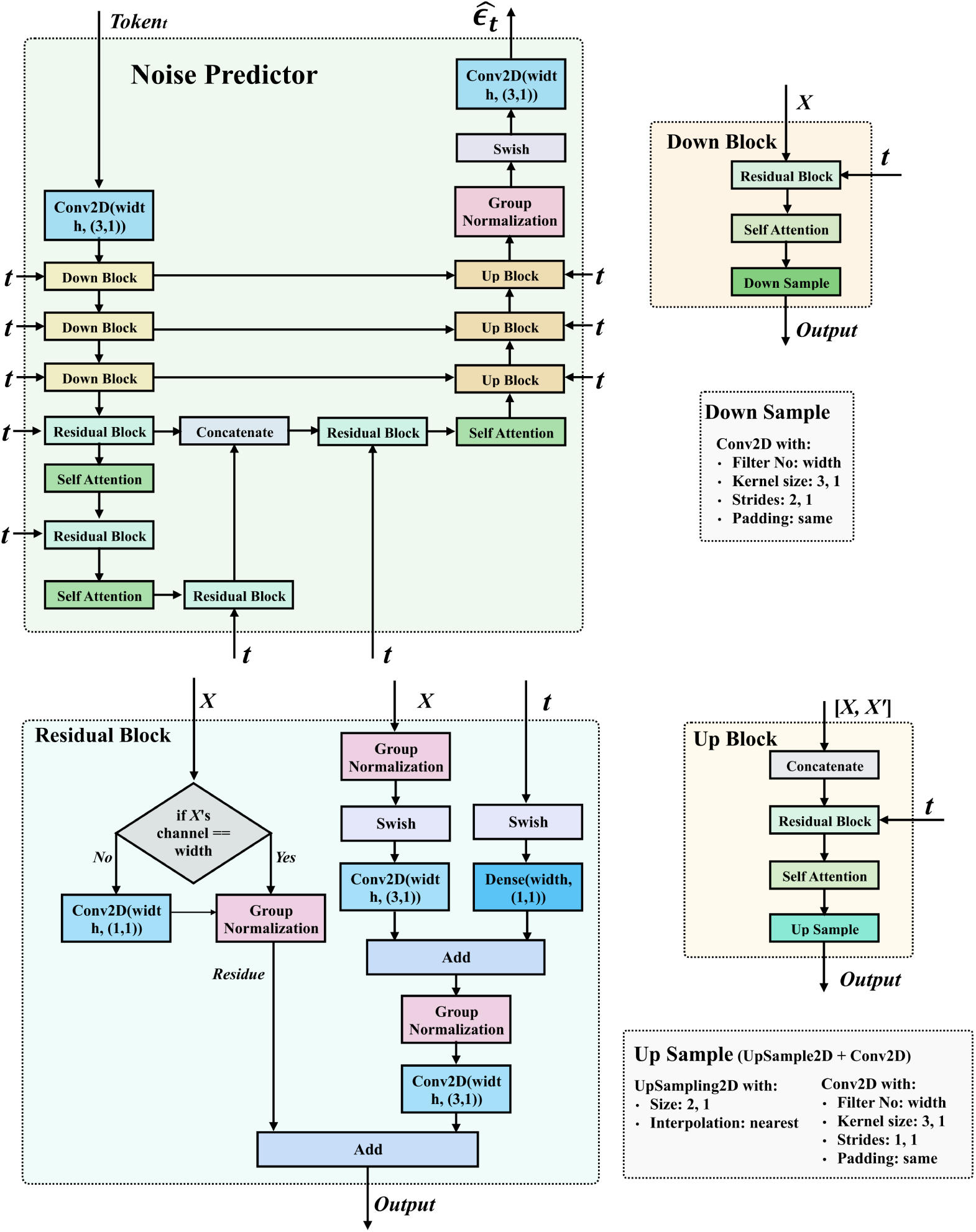
The noise predictor model. The upper left module shows the whole architecture of the noise predictor. The details of the residual block, the down block, and the up block of the noise predictor modules are listed in the bottom left, upper right, and bottom right parts, respectively. The two gray shadow blocks represent the parameters selected for the down sample and the up sample.

As shown in the upper middle part of Figure 2, the noisy classifier model sequentially consisted of the same architecture as the noise predictor model, a flatten layer, a dense layer with 32 neurons, and a dense layer with 1 neuron and sigmoid activation as outputs. where the output of *ŷ* is the AMP probability score, whose value is between 0 and 1. The larger the value of *ŷ* is, the higher the probability that the peptide has antimicrobial activity.

In the upper right box of Figure 2, the procedure that computes the *t* − 1 timestep classifier guidance noised token *ClsToken_t−_*_1_ from the *t* − 1 timestep noised token *Token_t−_*_1_. First, we extract classifier guidance from the input *Token_t−_*_1_ and the given guidance label *y*. Consequently, the guidance scale was utilized to multiply the classifier guidance and add *Token_t−_*_1_ to obtain the classifier guidance noisy token *ClsToken_t−_*_1_.

As shown in the bottom part of Figure 2, the *Token_T_*, also referred to as arbitrarily sampled Gaussian noise, is input into the denoise module to obtain the *Token_T−_*_1_ along with the given label *y* as input to the calculate classifier guidance *Token_t−_*_1_ module, and we obtain the *ClsToken_T−_*_1_. After *T* runs the denoise and the calculation classifier guidance *Token_t−_*_1_ modules, we obtain the generated clean token *ClsToken*_0_. Finally, we generate the candidate AMP sequences by unscaling *ClsToken*_0_ to 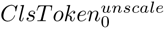 to *Sequence* on the basis of the token dictionary Eq. (16) and Eq. (2), respectively.

### Noise predictor

As shown in Figure 3, we use a U-Net as the architecture of the noise predictor model. The U-Net is constructed by several modules, including the residual block, down block, and up block, and some essential layers contain 2D convolutional layers, swish activation functions, and self-attention layers.

The residual block was first introduced as part of the ResNet architecture [41] to address the vanishing gradient problem, enabling the training of very deep neural networks by incorporating a skip connection part (the input to the residual block is added to its output). The process ensures that the network can learn not only the identity function (skip connection part) but also any other transformations (transformation part) to minimize the overall loss, which results in improving the ability to generalize and learn complex patterns. For the skip connection part of our residual block, the input hidden feature (*X*) is processed by a Conv2D function to enable the output’s channel number (*width*) to meet the network’s requirements. Then, group normalization is utilized to generate the identity as the output of the skip connection part. For the transformation part, the input hidden feature (*X*) is processed via group normalization, a swish activation function, and a Conv2D function. The timestep (*T*) is incorporated by a swish activation function and a dense layer (known as the fully connected layer) with a neural number of *width* and then adds the output and the former middle output of the transformation part together. Subsequently, group normalization and Conv2D are utilized to learn the transformation hidden features from the input hidden feature (*X*) and timestep (*T*) together. Finally, we add the outputs of the skip connection part and the transformation part together.

The down block aims to decrease the input matrix (*X*) to its half size via a residual block, a self-attention layer and a downsampling layer, which is a 2D convolutional layer with a filter number of *width*, kernel size of (3, 1), stride of (2, 1), and padding of ”same”.

The up block aims to increase the concatenate output of its input matrix (*X*), the output of its paired input matrix (*X^′^*), to its double size by a residual block, a self-attention layer and an upsampling layer. The upper sample layer consists of a 2D upsampling layer with a kernel size of (2, 1), an interpolation method of ”nearest”, a 2D convolutional layer with a filter number of *width*, a kernel size of (3, 1), a stride of (1, 1), and a padding of ”same”.

### Performance metrics

To evaluate the generative performance of the proposed model, generation accuracy (GenAcc) was computed. This accuracy is defined as the ratio of the number of peptides generated as positive by the proposed AMP classifier to the total number of valid generated sequences. The self-developed RFClassifier-AMP30 model was used to determine whether the generated sequences were AMPs. Validly generated valid sequences are those that consist solely of the 20 standard amino acids and have lengths between 5 and 30.

#### 2.2.4 AMP Classifier

### RFClassifier-AMP30

To comprehensively evaluate the generative performance of ClsDiff-AMP30, we developed an accurate AMP classifier to check the probability of whether the generated sequences are AMP or not. As shown in Table 2, we compared 14 traditional machine learning classifiers with 10-fold cross-validation (CV) using the AMP and non-AMP datasets. We then sorted all the results by accuracy and found that the RF classifier performed best. We then tune the hyperparameters of the RF algorithm via a grid search of ntree (*ntree* ∈ {100, 200*, …,* 2000}) and max features (”log2” or ”sqrt”) to find the best classifier model. Finally, we find that the RF with a ntree of 2000 and max features of ”log2” works best. The model is hereby named RFClassifier-AMP30.

**Table 2:**
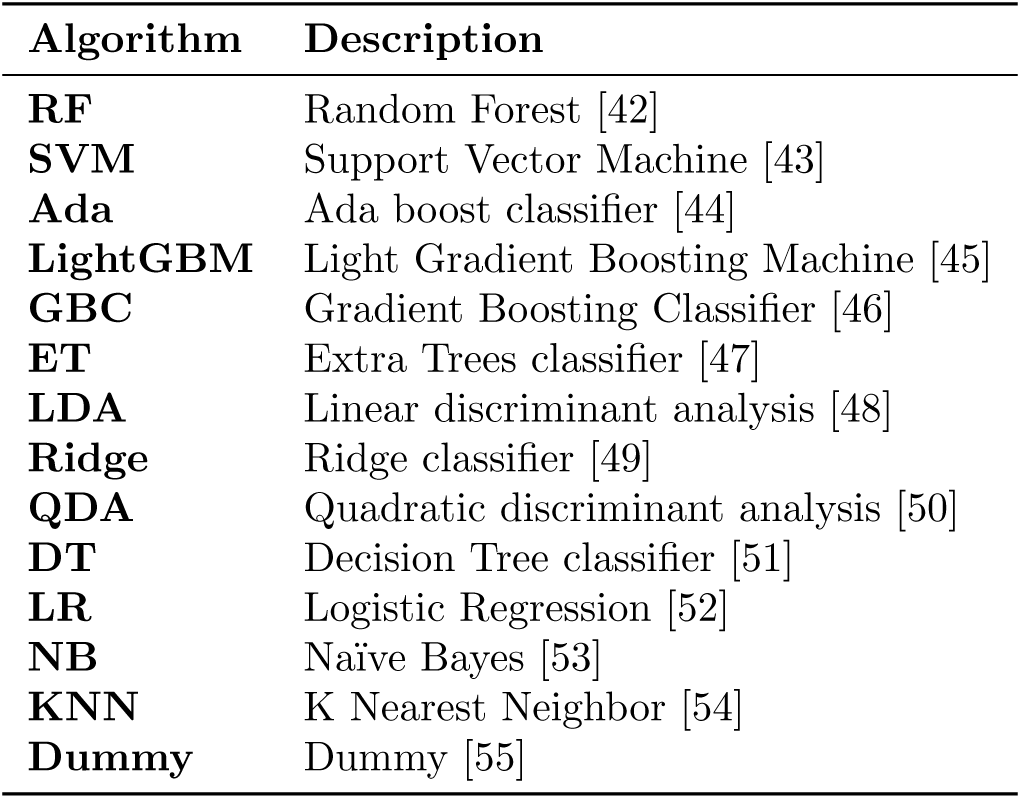
The fourteen traditional machine learning algorithms evaluated for the classification of AMP.

#### Input feature combinations

Following the classification and regression models of Yan et al. [39, 56], type3Braac9, type7raac19, composition of k-spaced amino acid pairs (CKSAAP) [57], dipeptide deviation from the expected mean (DDE) [58], and the k-spaced conjoint triad (KSCTriad) [59] were found to perform better than the other features.

These features were concatenated to train all the traditional machine learning classification models. Herein, type 3Braac9 and type 7raac19 belong to a collection of feature encoding methods called pseudo K-tuple reduced amino acid composition (PseKRAAC) [60]. PseKRAAC counts the number of reduced amino acid groups as the feature vector. PseKRAAC partitions all 20 natural amino acids into clusters on the basis of their different physical–chemical features. For example, type 3B and type 7 are two different types of amino acid clustering methods. In addition, each clustering method groups the 20 amino acids into *g* (*g* ∈ {1, 2, 3*…,* 20}) groups. The suffix in the name, e.g., raac*N*, means that the 20 amino acids are grouped into *N* reduced amino acid groups.

#### Performance metrics

We used the accuracy, area under the receiver operating characteristic (ROC) curve (AUC), recall, precision (Prec.), F1 score, kappa coefficient, and Matthews correlation coefficient (MCC) were used as performance metrics to evaluate the AMP classifier models.

#### 2.2.5 Wet Lab Experiments

##### MIC Determination

The minimal inhibitory concentration (MIC) of the AMPs was determined against *Escherichia coli* ATCC 25922, *Staphylococcus aureus* ATCC 25923, and methicillin-resistant *S. aureus* (MRSA, ATCC 43300) in accordance with the Clinical and Laboratory Standards Institute (CLSI) broth microdilution method M07–A11 [61]. The peptides were serially diluted twofold in Mueller–Hinton broth across a 96-well microtiter plate to achieve a concentration range of 256 to 0.5 *µ*g*/*mL in a 100 *µ*L volume. An equal volume (100 *µ*L) of bacterial suspension was subsequently added to each well, yielding a final inoculum concentration of approximately 5 × 10^5^ CFU/mL. The growth control contained Mueller–Hinton broth and bacteria without peptide, and the negative control contained Mueller–Hinton broth only. The plates were incubated at 37 *^◦^*C for 24 h, after which bacterial growth was assessed by measuring the optical density at 600 nm. The MIC was defined as the lowest peptide concentration that completely inhibited visible growth, equivalent to the negative control. All the assays were performed in triplicate.

### Hemolytic Activity Assay on Human RBCs

The percent hemolytic activity of the peptides was assessed against human red blood cells (RBCs) according to a protocol based on Sabo *et al.* [62]. Whole blood from healthy donors was centrifuged at 1,700×*g* for 5 min, and the resulting erythrocyte pellet was washed three times with phosphate-buffered saline (PBS; 137 mM NaCl, 10 mM Na_2_HPO_4_, 1.8 mM NaH_2_PO_4_, pH 7.4). A 1% (v/v) suspension of RBCs was prepared in PBS. The peptides were serially diluted twofold in PBS across a 96-well PCR plate to achieve a concentration range of 256 to 8 *µ*g*/*mL in a 50 *µ*L volume. An equal volume (50 *µ*L) of the 1% RBC suspension was added to each well, followed by incubation at 37 *^◦^*C for 1 h. PBS and 1% Triton X-100 served as negative (0% hemolysis) and positive (100% hemolysis) controls, respectively. After incubation, the plate was centrifuged at 1,700 × *g* for 5 min. Subsequently, 80 *µ*L of each supernatant was carefully transferred to a clear, flat-bottom 96-well plate, and hemoglobin release was quantified by measuring the absorbance at 405 nm. The percent hemolysis was calculated for each peptide concentration, and all the assays were performed in triplicate. The hemolysis percentage was calculated as follows:

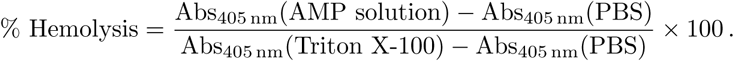

## 3 Experimental Results

The proposed ClsDiff-AMP30 model is highly versatile and capable of generating diverse peptide sequences and producing peptides of various sizes. We systematically evaluated its effectiveness and presented the results in five subsections. The first and second subsections describe the results of the hyperparameter search for RFClassifier-AMP30 and ClsDiff-AMP30, respectively. The third subsection evaluated the performance of the ClsDiff-AMP30 model in generating AMPs and provided a comprehensive analysis of the experimental findings. In the fourth subsection, we investigated the robustness of the model via an ablation experiment. Finally, we examined the 3D structures of the twelve new AMP candidates and analysed the sequence motifs of the 130 predicted AMPs.

### 3.1 Hyperparameter search for RFClassifier-AMP30

To evaluate the performance of the generative sequences, we sought to identify the optimal classifier for distinguishing AMPs from non-AMPs. To this end, fourteen traditional machine learning algorithms were evaluated via 10-fold CV with default parameters, and the classifiers were ranked on the basis of their validation accuracies. As shown in Table 3, the random forest classifier outperformed the other models in all the performance metrics except recall, achieving an accuracy of 0.890. Then, we conducted a grid search to optimize the hyperparameters of the random forest classifier. We explored ntree values from 100–2000 (in increments of 100) and max_features of log2 and sqrt via 10-fold CV. As shown in Figure 4, the optimal performance was achieved with ntree = 2000 and max_features = log2. Finally, the optimal model yields an accuracy of 0.901, an AUC of 0.958, an F1 of 0.905 and an MCC of 0.805.

**Figure 4:**
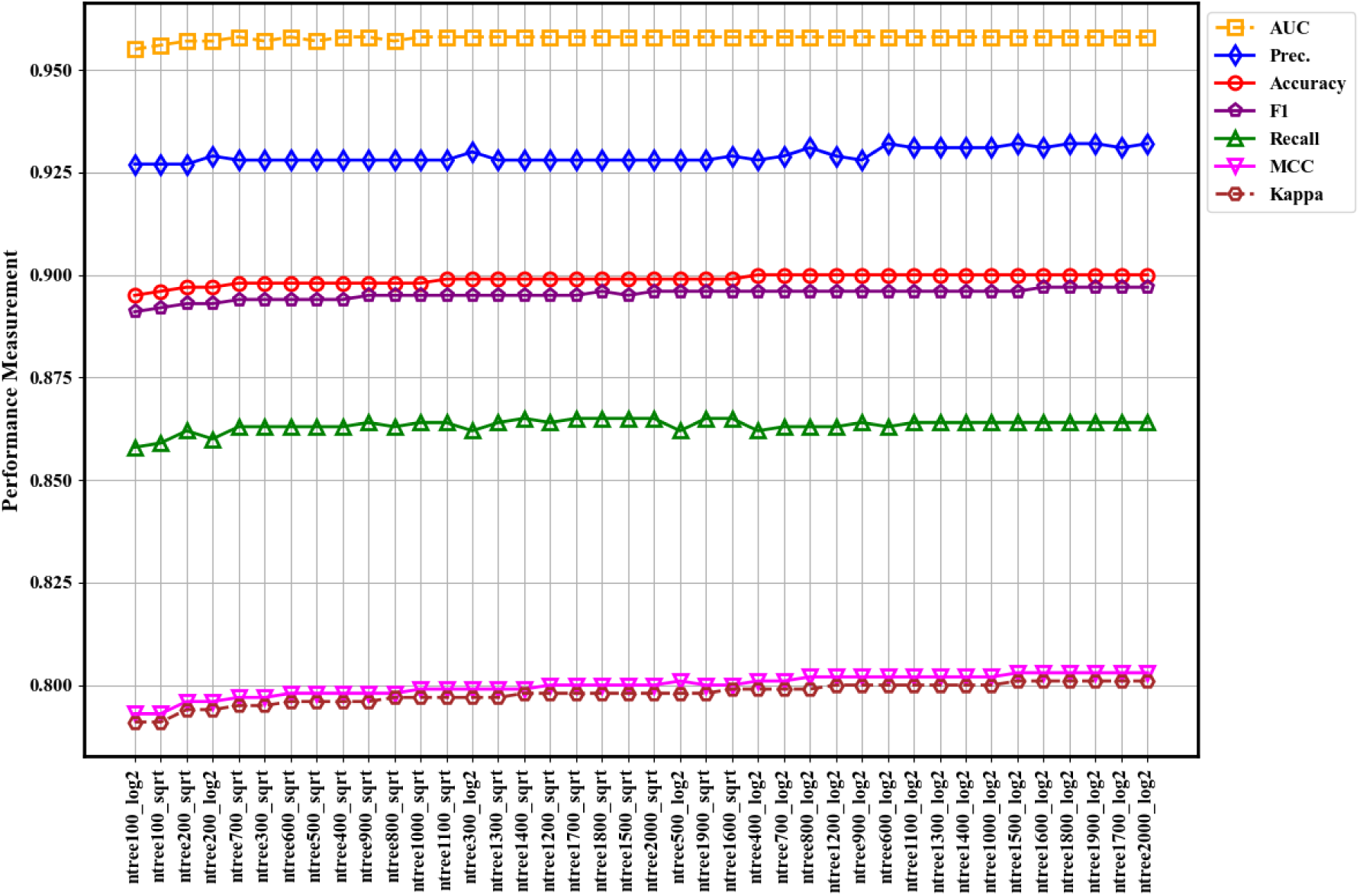
Grid search evaluation of ntree and max_features hyperparameters by 10-fold cross-validation in random forest models using AMP and non-AMP datasets.

**Table 3:** Tenfold CV classification performance of 14 traditional machine learning classifiers and the optimal model after grid search (RFClassifier-AMP30) on the AMP and non-AMP datasets.

| Model | Accuracy | AUC | Recall | Prec. | F1 | Kappa | MCC |
| --- | --- | --- | --- | --- | --- | --- | --- |
| **Random Forest Classifier (RFCClassifier-AMP30)<sup>†</sup>** | **0.901** | **0.958** | 0.937 | **0.875** | **0.905** | **0.803** | **0.805** |
| Random Forest Classifier | 0.890 | 0.955 | 0.840 | 0.934 | 0.884 | 0.780 | 0.784 |
| Extra Trees Classifier | 0.888 | 0.951 | 0.853 | 0.918 | 0.884 | 0.777 | 0.779 |
| Light Gradient Boosting Machine | 0.882 | 0.952 | 0.860 | 0.900 | 0.879 | 0.763 | 0.764 |
| Logistic Regression | 0.870 | 0.943 | 0.858 | 0.880 | 0.869 | 0.741 | 0.741 |
| Ridge Classifier | 0.870 | 0.000 | 0.855 | 0.882 | 0.868 | 0.740 | 0.741 |
| Linear discriminant analysis | 0.868 | 0.938 | 0.855 | 0.879 | 0.866 | 0.737 | 0.737 |
| SVM - Linear Kernel | 0.849 | 0.000 | 0.847 | 0.850 | 0.848 | 0.697 | 0.698 |
| Gradient Boosting Classifier | 0.839 | 0.920 | 0.773 | 0.890 | 0.828 | 0.678 | 0.684 |
| Naïve Bayes | 0.826 | 0.864 | 0.789 | 0.853 | 0.820 | 0.653 | 0.654 |
| Ada Boost Classifier | 0.811 | 0.893 | 0.775 | 0.835 | 0.804 | 0.622 | 0.623 |
| Decision Tree Classifier | 0.789 | 0.789 | 0.799 | 0.783 | 0.791 | 0.577 | 0.577 |
| K Neighbors Classifier | 0.669 | 0.839 | 0.955 | 0.608 | 0.743 | 0.339 | 0.413 |
| Quadratic discriminant analysis | 0.500 | 0.500 | **0.999** | 0.500 | 0.667 | 0.000 | 0.000 |
| Dummy Classifier | 0.500 | 0.500 | 0.400 | 0.200 | 0.267 | 0.000 | 0.000 |

<sup>†</sup> Model with optimal hyperparameters identified via a grid search

### 3.2 Hyperparameter search for ClsDiff-AMP30

We trained the proposed ClsDiff-AMP30 model using the AMP dataset, which contains 27,180 AMP sequences with lengths ranging from 5–30. The ClsDiff-AMP30 model consists of 2 submodels: the noise prediction model and the noise classifier model. They are integrated via a guidance scale parameter to generate sequences with elevated AMP scores. The amino acids were converted into the corresponding tokens as inputs, allowing the generation of novel AMP sequences from arbitrary Gaussian noise. During the training process, different training epochs were tested, and the generated sequences were evaluated via RFClassifier-AMP30 to assess the generative performance of the model. A total of 10,000 sequences were generated each time. The model with the optimal hyper-parameters, including the guidance scale, was systematically tuned via grid search in four stages:

*1. Initial configuration and epoch optimization of the noise predictor* We initialized the timestep parameter at 300, a value chosen to balance computational efficiency during preliminary training with flexibility for subsequent parameter refinement. We tested epoch values ranging from 100–10,000 and measured generation accuracy (GenAcc), defined as the proportion of generated sequences predicted to be AMPs by RFClassifier-AMP30. As shown in Figure 5(a), the model achieved peak performance at 7000 epochs, with a GenAcc of 55.98%. Thereafter, the performance plateaued, indicating that the model capacity had developed sufficiently through extended training iterations.
*2. Timestep refinement of noise predictor* With the epoch parameter fixed at 7000, we systematically optimized the timestep values ranging from 100–1000. As shown in Figure 5(b), the model achieves an improved GenAcc of 59.01% with a timestep of 700. A longer timestep enables greater precision in the diffusion process, which leads to improved precision in the generation of AMPs.
*3. Epoch optimization of the noisy classifier and best guidance scale exploration* To steer the generation toward realistic AMPs, the classifier guidance approach was introduced using the noised classifier. We systematically tested the number of epochs for the classifier from 100–10000. Figure 6 shows the performance of the model in terms of GenAcc and other metrics. The best performance was achieved at 3000 epochs, with an accuracy of 81.00%. This significant improvement confirms the effectiveness of the classifier guidance approach for AMP generation.
*4. Guidance scale calibration* The final optimization of the scale hyperparameter with a fixed number of 700 timesteps and 3000 epochs identified 75.0 as the optimal scaling factor. The resulting 65.76% accuracy shown in Figure 5(c) demonstrates the scale’s critical function in modulating classifier guidance effectiveness during generation.

**Figure 5:**
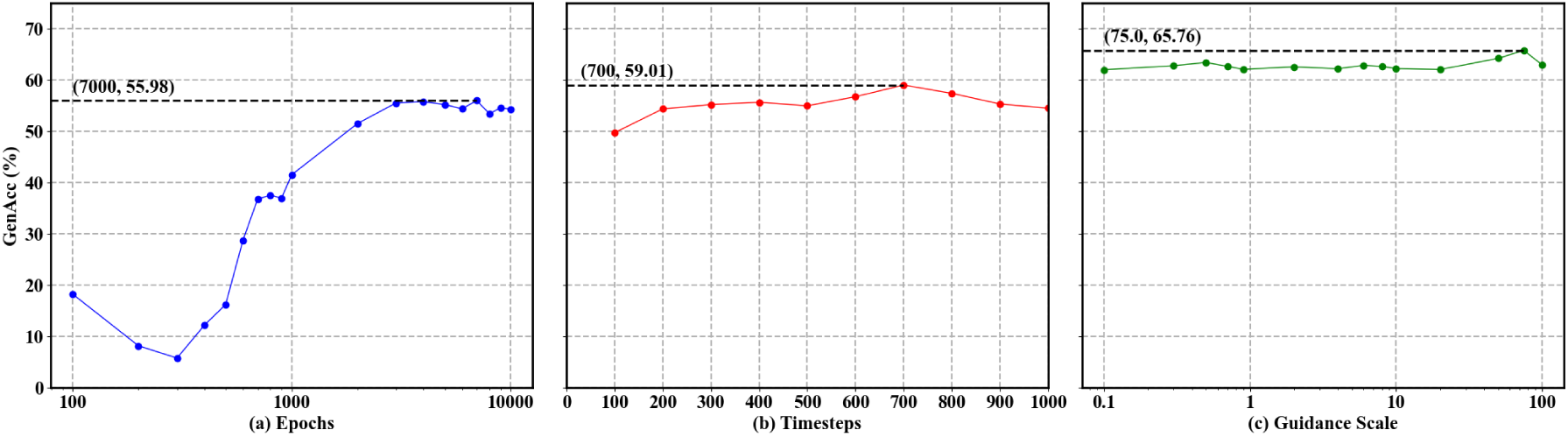
Hyperparameter optimization for ClsDiff-AMP30 using the AMP dataset. The figures (from left to right) illustrate the change in GenACC for the 10,000 generated peptides at different (a) training epochs, (b) diffusion timesteps, and (c) classifier guidance scales.

**Figure 6:**
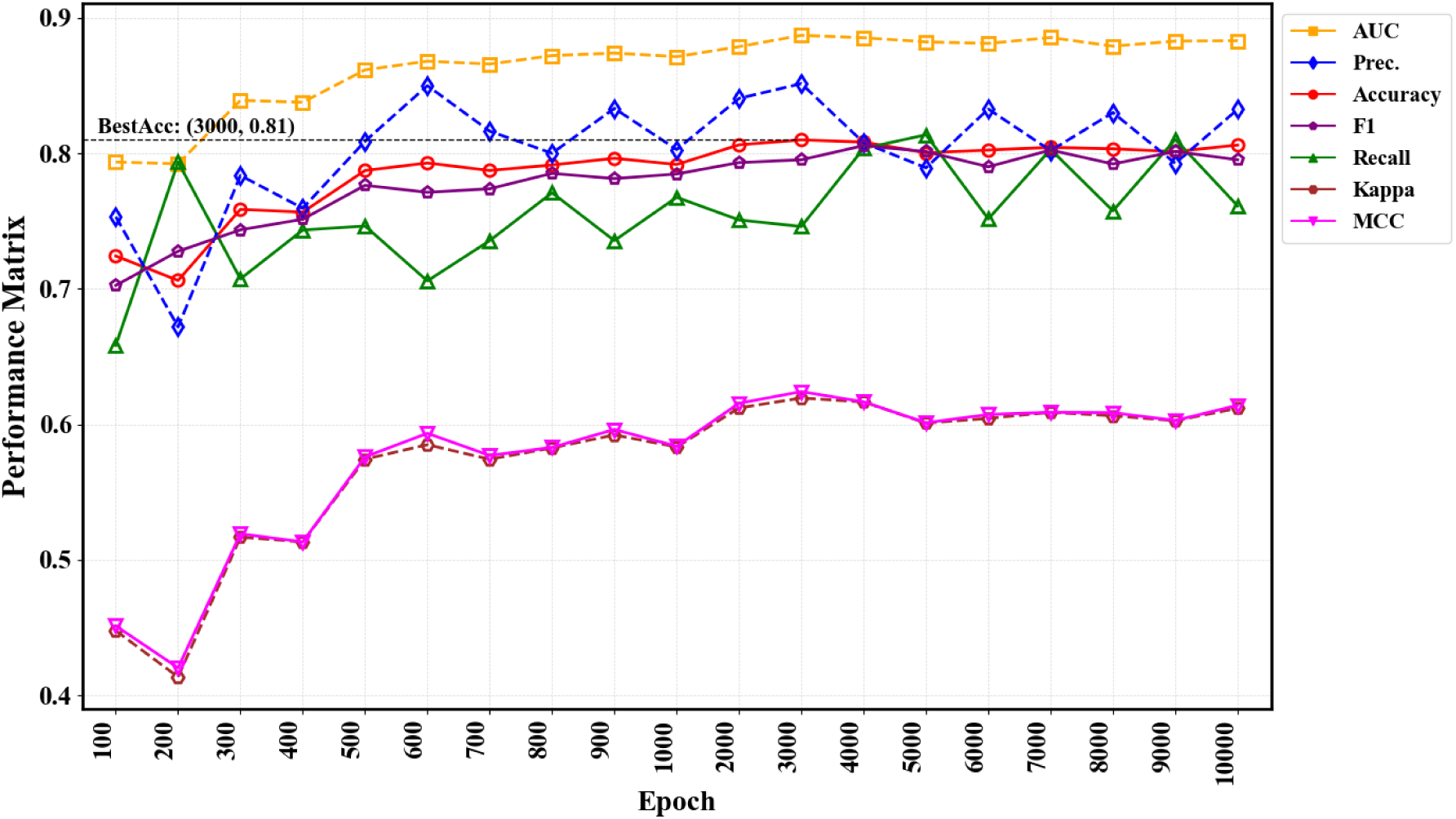
Comparative grid search evaluation of various epochs through Noised Classifier Models, using AMP and non-AMP datasets.

To summarize, we determined the optimal combination of hyperparameters for the ClsDiff-AMP30 model through systematic grid searches. The optimal values are as follows: Epochs are 7000 and 3000 for the noise predictor and the noise classifier, respectively. The diffusion timestep is 700, and the guidance scale is 75. This configuration significantly improved both the accuracy and stability of the ClsDiff-AMP30 model, providing a robust foundation for subsequent experiments.

### 3.3 Sequence and physicochemical characterization of generated AMPs

To validate the generative capability of the proposed ClsDiff-AMP30 model, a series of quantitative analyses were performed to compare the generated AMPs with real AMPs, randomly generated sequences, and randomly selected peptides from natural sources. Here, we define the four sets of sequences for the comparative study:

- GeneAMP: The sequences generated by ClsDiff-AMP30 and predicted with AMP scores greater than 0.5, i.e., classified as AMPs. Using the optimal model, we generated 10,000 sequences, of which 6,576 sequences were classified as AMPs.
- RealAMP: The sequences were randomly selected from the AMP databases.
- RndPep: The sequences were randomly selected from the PeptideAtlas database.
- RndSeq: The sequences were randomly generated with a uniform length distribution and uniform amino acid composition.

For comparative study purposes, each of the RealAMP, RndSeq, and RndPep sets contains the same number of sequences as GeneAMP, which is 6,576 sequences.

Furthermore, four analytical experiments were conducted. The first is the length distribution comparison of the four datasets. The second and third experiments are principal component analysis (PCA) scatter plots based on the amino acid composition (AAC), and the combined feature groups include ”Type3Braac9”, ”Type7raac19”, ”CKSAAP”, ”DDE”, and ”KSCTriad”. Finally, we generated a PCA scatter plot for eight physicochemical properties, including charge, charge density, isoelectric point, instability index, aromaticity, aliphatic index, boman index, and hydrophobicity ratio, on the basis of the four datasets. We also constructed eight violin plots to visualize the distributions of the eight physicochemical properties in the four datasets.

#### 3.3.1 Comparison of sequence length distributions

In this section, we analysed GeneAMP, RealAMP, RndPep, and RndSeq, focusing on sequence length distributions and amino acid compositions. As illustrated in Figure 7, the distributions of sequence lengths of GeneAMP and RealAMP are similar, both having irregular shapes with multiple peaks. In contrast, RndPep has a different shape with a broad distribution towards shorter lengths (9–16 residues), whereas RndSeq, which is drawn from a uniform distribution, has less variability and thus a uniform shape in the distribution. This result indicates that the set of generated and positively predicted AMPs may share the length characteristics of real sequences.

**Figure 7:**
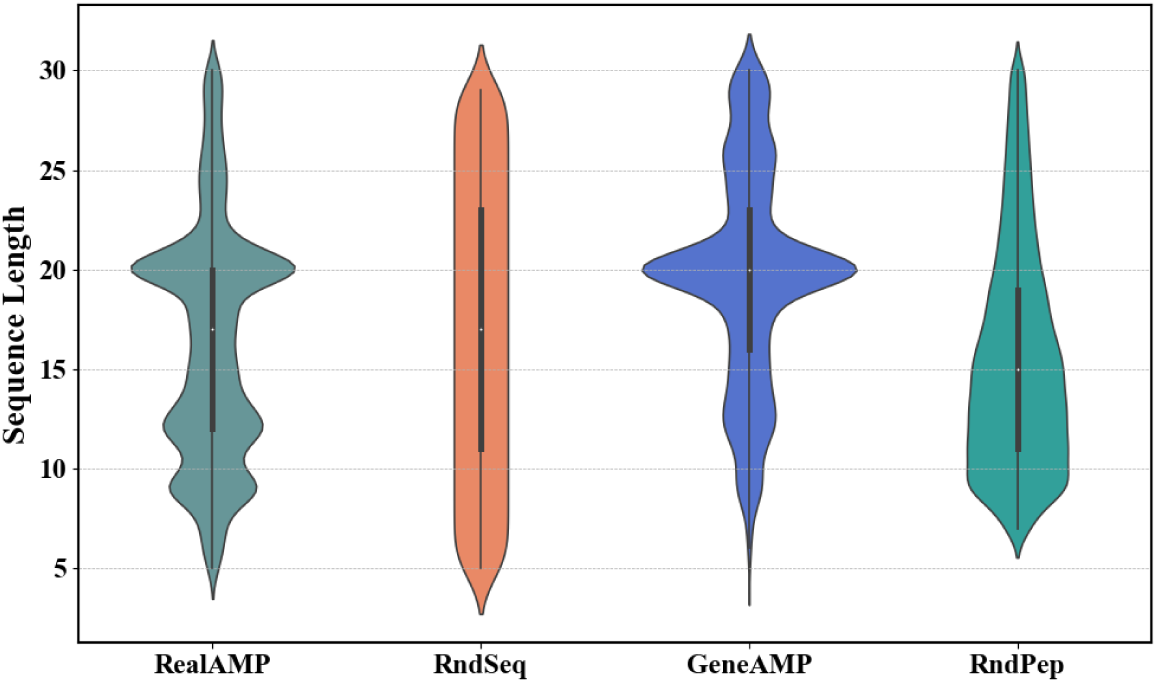
Comparison of the distribution of sequence lengths in the RealAMP, RndSeq, GeneAMP, and RndPep sets.

#### 3.3.2 Principal component analysis (PCA) of amino acid compositions

To evaluate how the generated sequences relate to the real and random peptide sets in terms of their physicochemical properties, we performed a comparative study of the amino acid composition (AAC) of the sequences. After computing the AAC features for each set of sequences, we applied principal component analysis (PCA) for dimensionality reduction to visualize the underlying distribution patterns of the set. The resulting scatter plots reveal both shared and distinct compositional characteristics in the four sequence sets.

As shown in Figure 8, GeneAMP has an uneven spread in the AAC feature space, which shows a higher degree of overlap with that of RealAMP. The data points range from approximately -3.0–4.0 in PCA1 and from approximately -4.0–4.0 in PCA2. The RealAMP data points could cover the range of the GeneAMP data points and even extend wider (PCA1: approximately -4.0–5.0; PCA2: approximately -4.0–4.5), suggesting that sequences in RealAMP are more diverse in their properties. On the other hand, the RndPep and RndSeq data points are clustered around the origin, indicating that there are no significant preferences or trends in the sequence sets.

**Figure 8:**
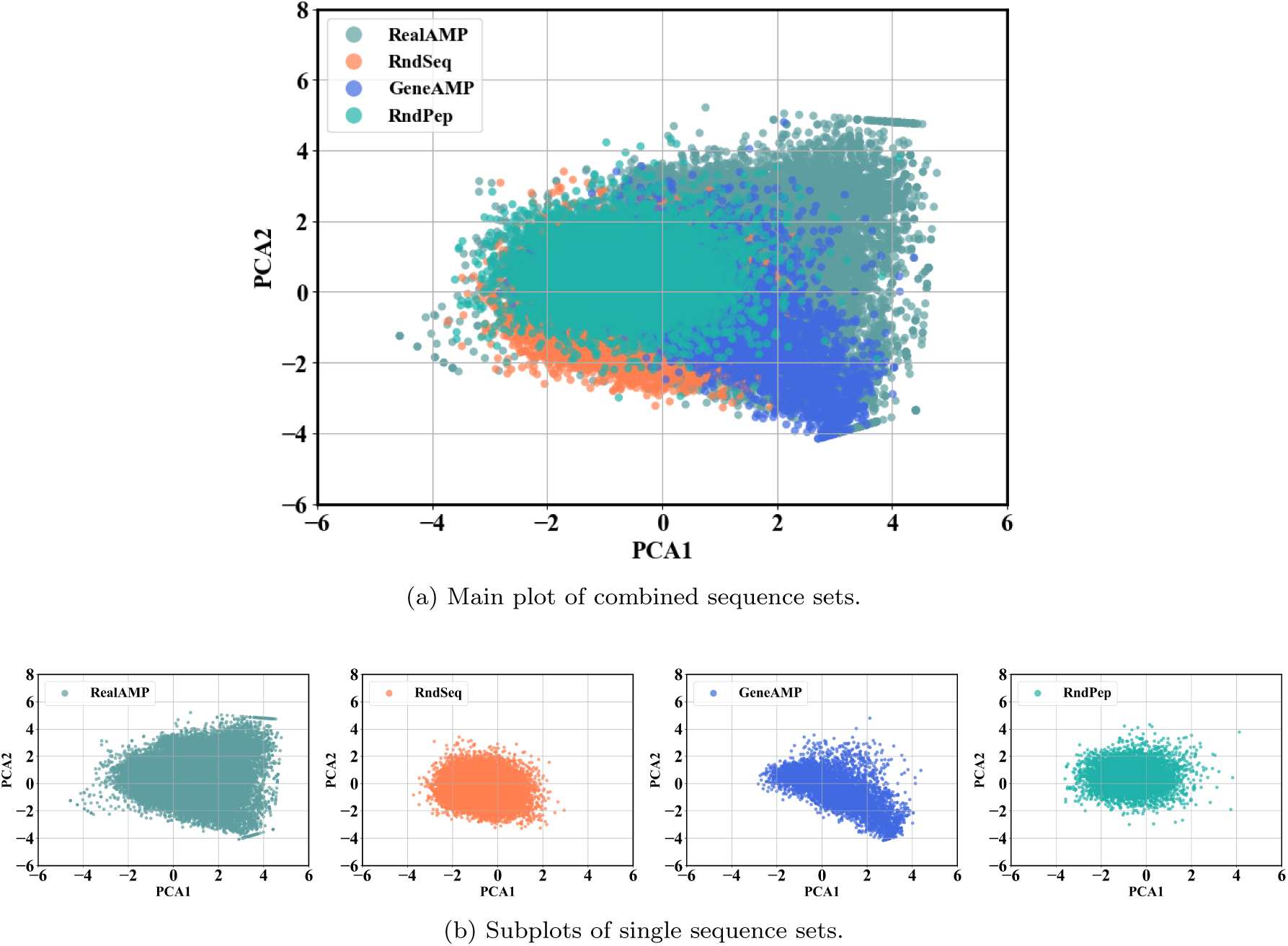
PCA scatter plots depicting the amino acid compositions of RealAMP, RndSeq, GeneAMP, and RndPep. Panel (a) displays the main plot with all the sets combined, whereas Panel (b) provides subplots for each sequence set.

In contrast, randomly generated sequences (RndSeq) show a distinctly different distribution, characterized by highly uniform and symmetric dispersion around the center, spanning approximately PCA1: -3.0–2.0 and PCA2: -3.0–2.5.

Overall, PCA visualization clearly illustrates the closer resemblance between GeneAMP and RealAMP in terms of amino acid compositional patterns, distinctly separating them from random distributions.

#### 3.3.3 Principal component analysis based on a combination of sequence features

To represent the sequences more comprehensively, we selected Five predictive amino acid encoding features, ”type3Braac9”, ”type7raac19”, ”CKSAAP”, ”DDE”, and ”KSCTriad”, are used to describe the sequences in GeneAMP, RealAMP, RndPep, and RndSeq. The same PCA was performed, and the results are shown in Figure 9.

**Figure 9:**
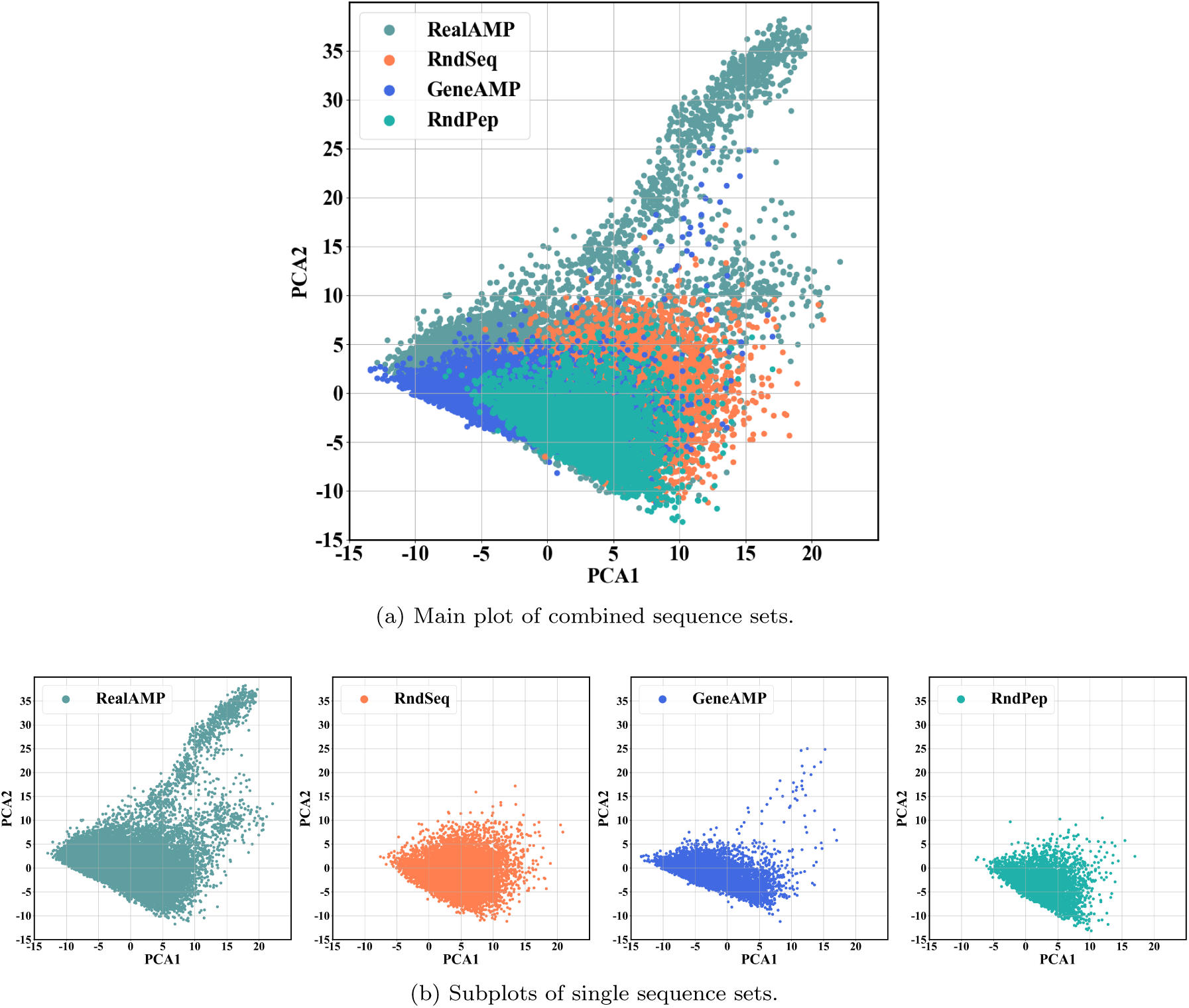
The PCA scatter plots illustrate the feature combinations (including ”type3Braac9,” ”type7raac19,” ”CKSAAP,” ”DDE,” and ”KSCTriad”) for RealAMP, RndSeq, GeneAMP, and Rnd-Pep. Panel (a) displays the main plots, whereas Panel (b) provides the subplots.

Notably, the shape and spatial distribution of the GeneAMP sequences are highly similar to those of the RealAMP sequences. Both sequence sets form a hat-like structure with two distinct ”ears” – one longer and one shorter – oriented toward the upper-right quadrant. Specifically, most GeneAMP and RealAMP sequences span the region of -15–15 along PCA1 and -10–25 along PCA2. While RealAMP covers a similar region, it extends beyond this range, spanning from -15–20 along PCA1 and -10–40 along PCA2.

In contrast, the RndSeq and RndPep sequences have similarly compact distributions, forming a smaller hat-shaped cluster. Most RndSeq sequences are located within -7–20 along PCA1 and -10–10 along PCA2, whereas RndPep sequences predominantly occupy -7–15 along PCA1 and -12–8 along PCA2.

#### 3.3.4 Principal component analysis on the basis of physicochemical characteristics

PCA was performed on the physicochemical features of the sequence sets, including charge, charge density, isoelectric point, instability index, aromaticity, aliphatic index, Boman index, and hydrophobicity ratio. The resulting scatter plots in Figure 10 reveal distinct clustering patterns among real, generated, random peptides, and random sequences.

**Figure 10:**
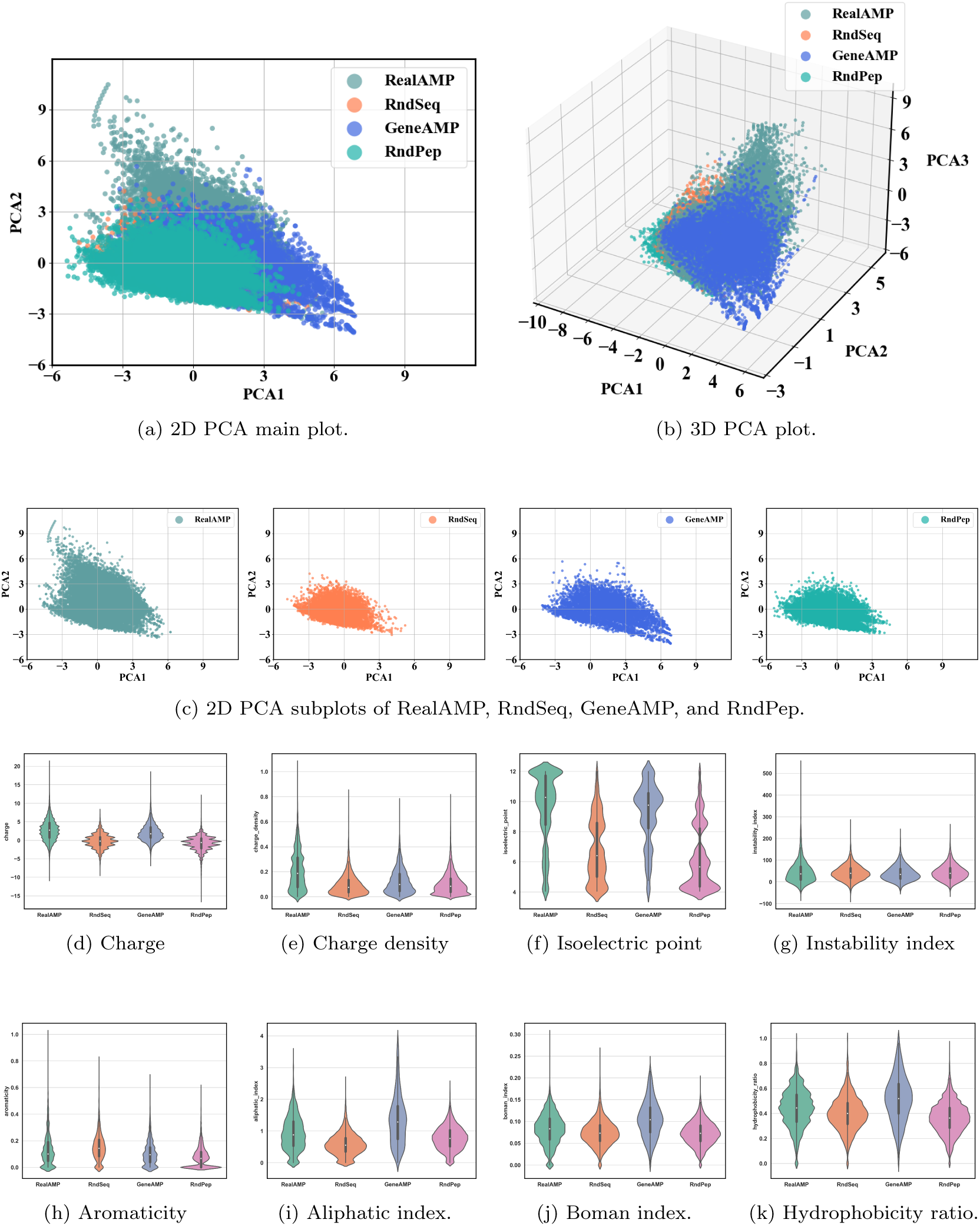
Physicochemical feature comparison: Panels (a) and (b) present scatter plots of sequences after extraction of physicochemical properties, including charge, charge density, isoelectric point, instability index, aromaticity, aliphatic index, boman index, and hydrophobicity ratio, followed by PCA for dimensionality reduction. Panel (c) shows the subplots, whereas the remaining panels display violin plots for each individual feature.

The GeneAMP sequences show a high degree of overlap with those in RealAMP. Figure 10 (c) shows that the GeneAMP sequences cluster mainly from -4.5–7 along PCA1 and from -5–4 along PCA2, which significantly overlaps with the RealAMP distribution range (PCA1: -4.5–5, PCA2: -3–9). This spatial proximity indicates that the model has learned and reproduced the composition and physicochemical properties of real AMP, with high fidelity in generation.

The RndSeq sequences show a much more dispersed distribution in the PCA space, as depicted in Figure 10 (c). These sequences span a wide range from -4.5–4 along PCA1 and -3–3 along PCA2, indicating a lack of compositional structure. The widespread of points underscores the randomness in the sequences, with no clear biological pattern or clustering. However, the RndPep sequences show a more concentrated distribution, mostly within -5–4 along PCA1 and -3–3 along PCA2. Although more structured than random sequences, RndPep sequences remain distinct from the RealAMP and GeneAMP clusters, indicating that they are still relatively less biologically coherent but more structured than completely random sequences.

In contrast, the RndSeq sequences are widely dispersed across the PCA space, lacking the compactness observed in the GeneAMP and RealAMP groups. This broad spread highlights the absence of biologically meaningful structures in random sequences, which are characterized by a lack of physicochemical consistency. Although RndPep sequences show a more concentrated distribution than RndSeq sequences do, they still exhibit a less structured nature than do the GeneAMP and RealAMP sequences Figure 10 (c) clearly demonstrates the similarity between the GeneAMP and RealAMP sequences, validating the model’s ability to generate biologically relevant peptides. In contrast, random sequences (RndSeq and RndPep) display a broader, less structured distribution, reinforcing the biological plausibility of GeneAMP sequences, which closely resemble real AMPs in both structure and physicochemical properties.

The above analyses collectively demonstrate that the GeneAMP sequences generated by the model closely resemble the RealAMP sequences, exhibiting strong overlaps in compositional patterns and physicochemical features. Specifically, both groups consistently cluster in similar PCA regions, highlighting the effectiveness of the model in capturing biologically relevant characteristics of authentic antimicrobial peptides. In contrast, the RndSeq and RndPep sequences presented dispersed PCA distributions and inconsistent physicochemical profiles. These findings validate the model’s ability to generate structured, biologically plausible AMP sequences rather than random or nonspecific peptides.

In addition, we examined the underlying physicochemical properties of the sequences. The violin plots of key parameters provide further evidence of the model’s ability to generate peptides whose characteristics closely resemble those of authentic AMPs, distinguishing them from random constructs. As shown in Figure 10 (d–k), except for the charge density and instability index, the GeneAMP sequences exhibit a high degree of similarity with RealAMP, clearly differentiating them from RndSeq and RndPep.

### 3.4 Validation of model robustness and ablation experiments

To comprehensively evaluate the robustness and generalizability of the proposed generative framework, we conducted a series of experiments under various conditions and settings. These include analyses of model performance across different noise distributions, the effectiveness of classifier-guided sampling, and generative performance under different data sizes. By systematically comparing variants of the model, we aim to validate not only its capacity to generate biologically meaningful peptide sequences but also the importance of design choices that contribute to its overall performance.

#### 3.4.1 Effect of noise distributions

To evaluate the impact of different noise priors on generative performance, we compared Gaussian, uniform, and gamma noise within our model framework. As shown in Table 4, the Gaussian distribution achieved the highest generation quality, producing 10,000 valid sequences with an accuracy of 65.76% under classifier guidance. In contrast, both uniform and gamma noise resulted in significantly lower accuracy and fewer valid sequences, even when classifier guidance was applied.

**Table 4:** Accuracy of the 10000 generated sequences under different noise distributions.

| Noise Distribution | Valid Sequences | GenAcc (%) |
| --- | --- | --- |
| Gaussian | 10,000 | 65.76 |
| Uniform | 8,624 | 8.38 |
| Gamma | 6,060 | 9.14 |

These results demonstrate that Gaussian noise provides a more stable and effective foundation for the denoising process in our model. Moreover, the accuracy improvement under classifier guidance highlights the positive contribution of our guidance module.

#### 3.4.2 Effect of dataset size

Figure 11 presents the impact of training dataset size on the performance of the noise predictor model, both with and without classifier guidance. As the size of the dataset increases, the generation accuracy consistently improves, particularly when the classifier guidance mechanism is applied. This trend confirms that our model effectively benefits from larger training datasets, enabling it to better learn sequence patterns and biological properties. Moreover, the consistent performance gain observed with classifier guidance across all dataset sizes further highlights the robustness and effectiveness of the guidance module in enhancing generation quality.

**Figure 11:**
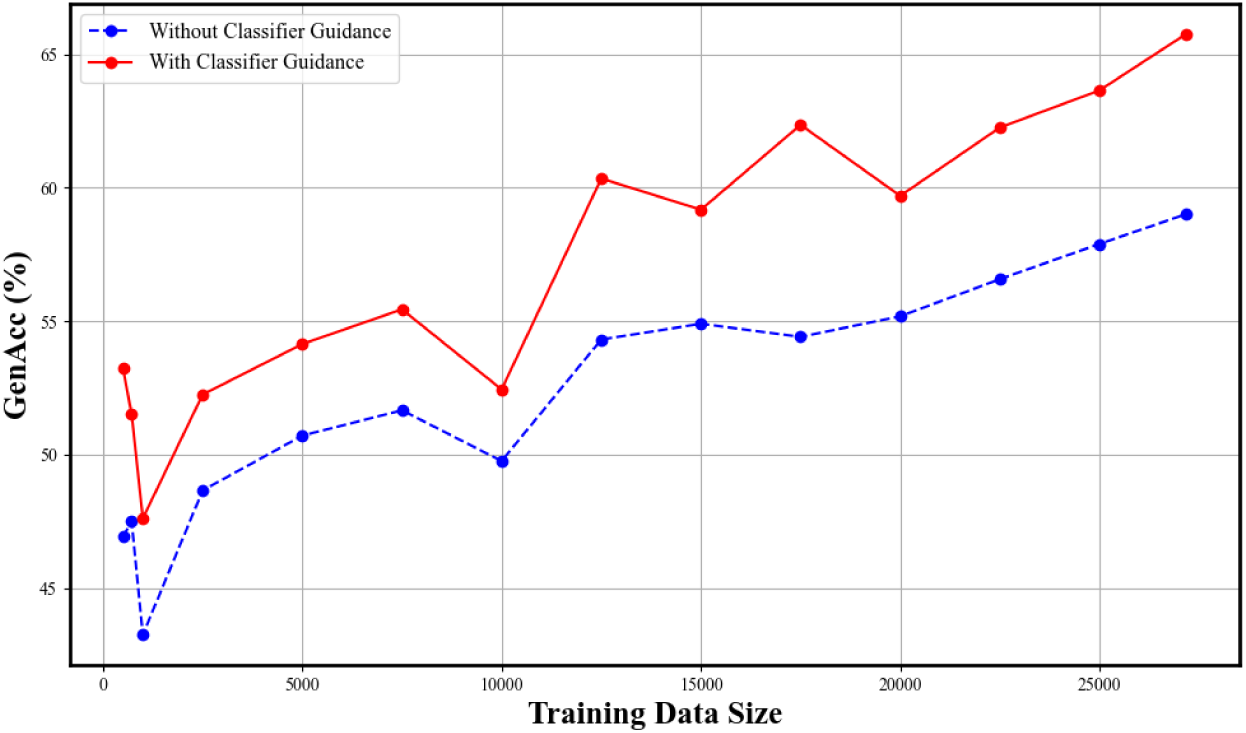
Accuracy of the generated sequences under different training data sizes.

#### 3.4.3 Effect of classifier guidance

To evaluate the effectiveness of classifier guidance for valid AMP generation, an ablation experiment was conducted to compare the results generated by the proposed model and the variant model with the classifier guidance component removed. Both models were trained under the same settings of 7000 epochs and 700 timesteps to ensure a fair comparison.

Table 5 shows a clear performance improvement when classifier guidance was employed. The unguided noise predictor achieved an accuracy of only 59.0%, whereas the guided noise predictor reached a significantly higher accuracy of 65.8%, when a guidance scale of 75.0 was used. This 11.5% improvement highlights the effectiveness of classifier guidance in steering the generation process toward peptides with enhanced antimicrobial actvitiy prediction. This finding demonstrates that classifier guidance improves the overall quality of the generated peptides.

**Table 5:** Ablation experiment comparing model performance with and without classifier guidance.

| Model | Valid Sequences | GenAcc (%) | Improvement |
| --- | --- | --- | --- |
| Without classifier guidance | 9999 | 59.0% | - - |
| With classifier guidance | 10000 | 65.8 | 11.5 |

### 3.5 Further analysis

#### 3.5.1 Sequence Simulation

To improve the experimental results and further validate the authenticity of the generated sequences, we conducted minimum inhibition concentration (MIC) prediction of the generated sequences against *Escherichia coli* (EC) and *Staphylococcus aureus* (SA) and 50% hemolytic concentration (HC_50_) prediction. The two MIC values of EC*_MIC_* and SA*_MIC_* are predicted via BERT-AmPEP60 [63]. HC_50_ values predicted by BERT-HemoPep60. After that, we screen 12 sequences that meet all the requirements in Eq. (17).

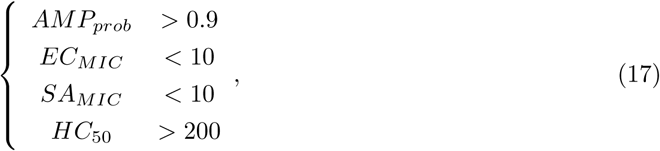

where AMP*_P rob_* indicates the predicted probability of being an AMP. In addition, EC*_MIC_*, SA*_MIC_*, and HC_50_ use the same unit of *µM* .

As shown in Table 6, a total of twelve sequences are selected, followed by Eq. 17. Consequently, we conducted a 3D structure simulation with PEP-FOLD3 [64]. Molecular dynamics simulations will be performed to analyse the interactions between AMPs and bacterial membranes. This approach enables high-dimensional analysis, providing insight into the biological relevance and functional integrity of the generated sequences while mitigating the potential loss of key features during dimensionality reduction. Table 6 presents the 12 selected sequences after filtering, and Figure 12 shows the corresponding 3D structures constructed via PEP-FOLD3. The 3D structure of most selected GenAMP is made of a single *α*-helixes, except for Seq7 which is formed of two short *α*-helixes linked by a loop and Seq5 and 10, which are random coils.

**Figure 12:**
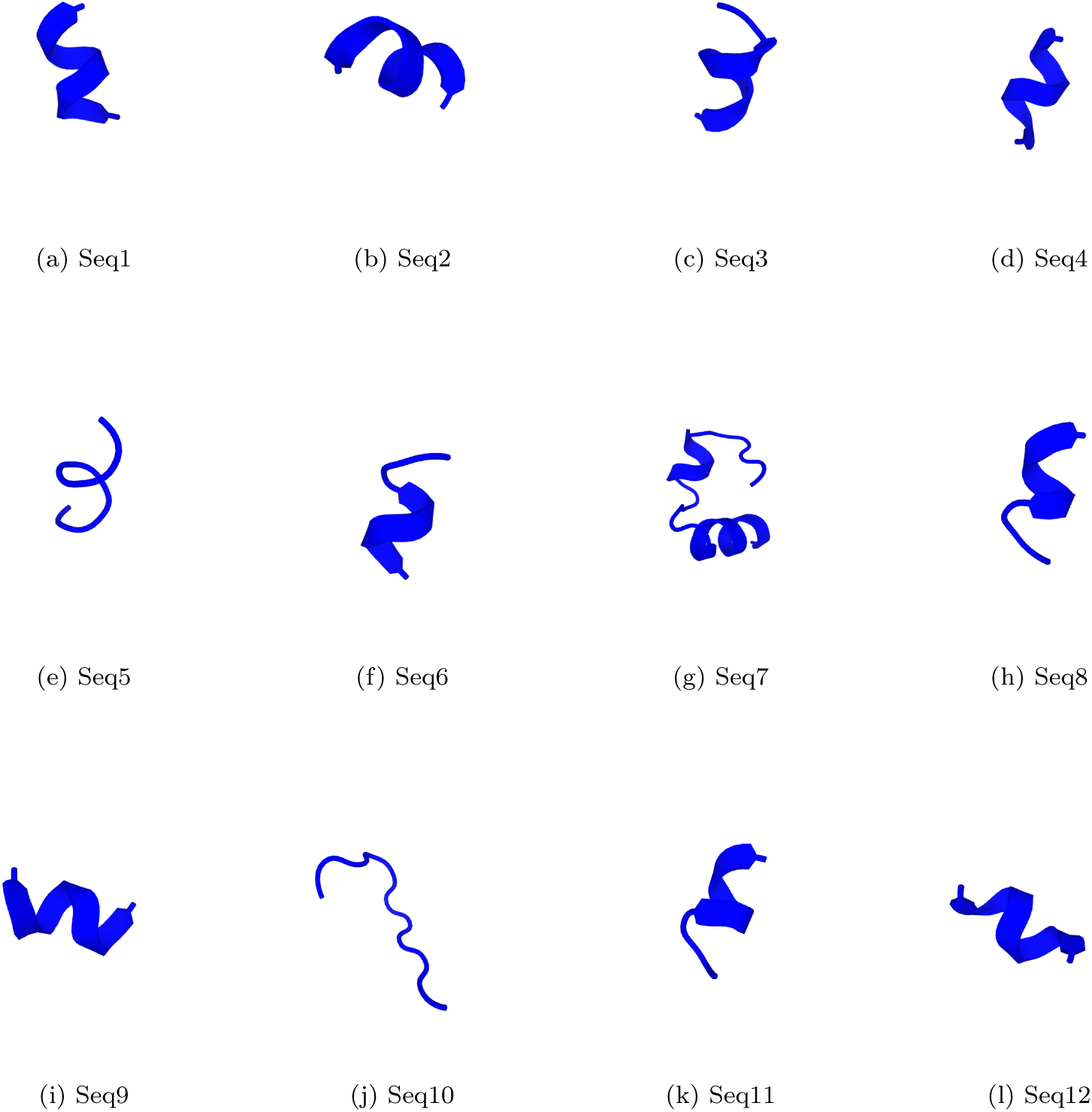
Sequence 3D structures generated by PEP-FOLD3.

**Table 6:** Filtered sequences with predicted antimicrobial activity and toxicity scores. Note: All MIC values for EC and SA, and HC_50_ are reported in *µM* .

| ID | Sequence | $AMP_{Prob} \uparrow^a$ | $HC_{50} \uparrow$ | $EC_{MIC} \downarrow^b$ | $SA_{MIC} \downarrow$ |
| --- | --- | --- | --- | --- | --- |
| Seq1 | RRWWRWWRI | **0.9975** | 247.93 | 5.36 | 3.72 |
| Seq2 | YRWWRWWRI | 0.9895 | 204.39 | 8.73 | 6.17 |
| Seq3 | WRTRRRWRY | 0.9825 | 313.96 | 8.16 | 3.11 |
| Seq4 | KRRWKTWWW | 0.9775 | 293.83 | 4.13 | 1.24 |
| Seq5 | RWKWRWRSW | 0.9770 | 225.01 | 8.35 | 8.16 |
| Seq6 | KWRRFKWWL | 0.9770 | 296.82 | 9.73 | 1.19 |
| Seq7 | KALTKLFKKIGIGFKKAVHVGKVALS | 0.9765 | 243.64 | 5.82 | 8.33 |
| Seq8 | RWRRWKYWY | 0.975 | **459.38** | **3.04** | 1.36 |
| Seq9 | KRRYRWIWI | 0.9745 | 214.31 | 7.47 | 4.41 |
| Seq10 | FRWRIVVIRYRR | 0.971 | 237.37 | 5.84 | **1.02** |
| Seq11 | LWRRWYRWI | 0.9700 | 304.91 | 9.94 | 6.43 |
| Seq12 | YKKRWRLWI | 0.9655 | 263.51 | 6.43 | 4.59 |

<sup>a</sup>Upwards arrow ( ↑ ): A higher metric value corresponds to better sequence performance.
<sup>b</sup>Downwards arrow ( ↓ ): signifies the opposite.

#### 3.5.2 Motif exploration

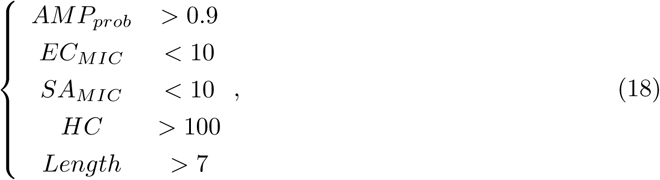

To explore the same patterns of generated AMP sequences, we generated several motifs and presented them via protein LOGO images via the Multiple EM for Motif Elicitation (MEME) suite [65]. For that purpose, we selected 130 sequences by following the criteria of Eq. (18). As shown in Eq. (18), we extend the threshold of HC from 200 of Eq. (17) to 100 to screen more sequences, find more and high-quality motifs, and add a length threshold of larger than 7 due to the same sequence length requirement of the MEME suite.

For Motif 1 (”GFCWRVCVYRNGVRVCHRRCN”) shown in Figure 13 (a), the *C* residues at positions 7, 16, and 20 and *R* residues at positions 14, 18, and 19 and *G* at position 12 are not recommended for modification. Position 1 strongly favours *G* but permits substitution with *W* or *Q*. Position 17 shows equal preference for *H*, *G*, *Y*, and *W*.

**Figure 13:**
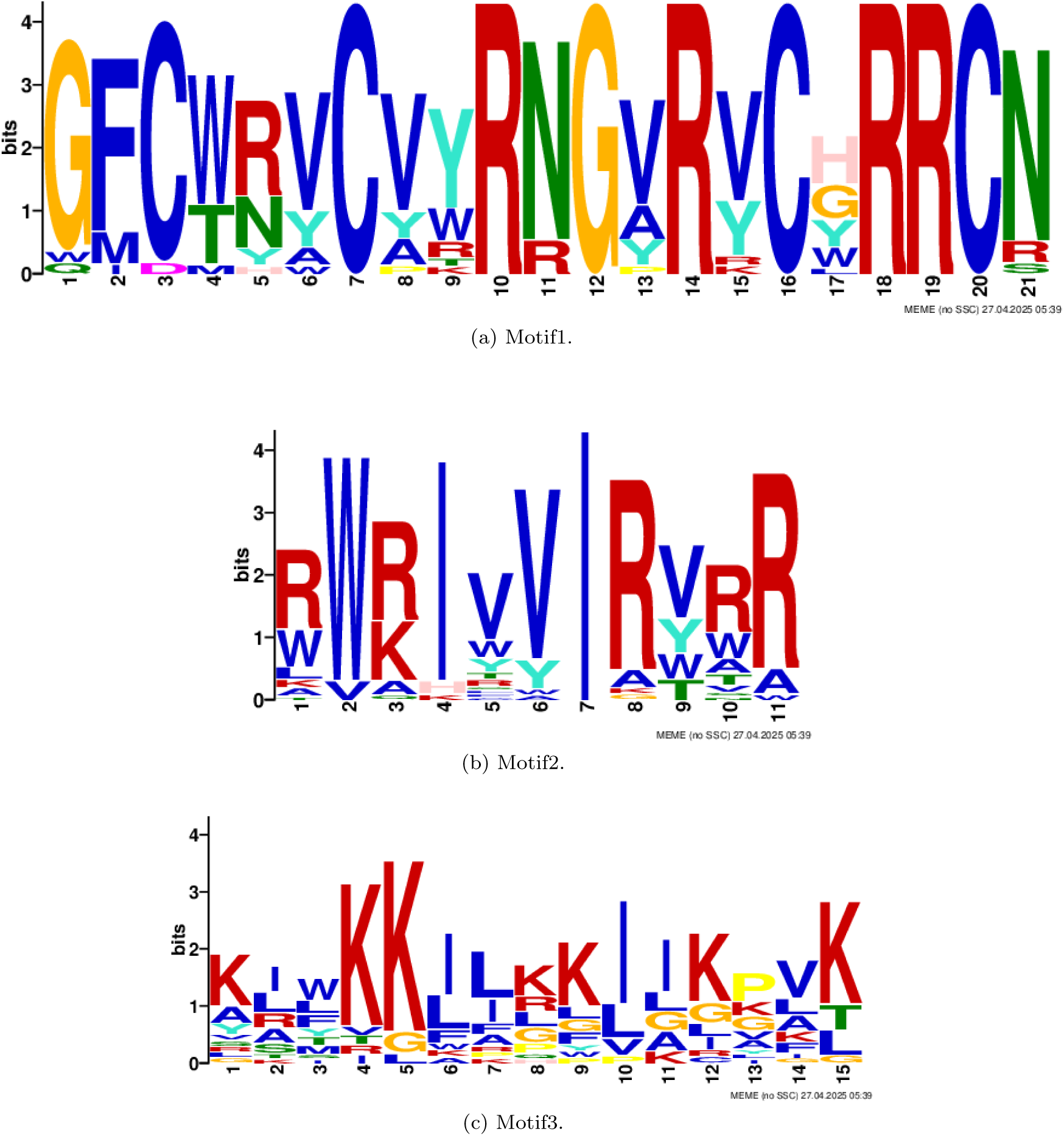
AMP motifs of the selected 130 generated sequences screened via Eq. (18).

Motif 2 (”RWRIVVIRVRR”) in Figure 13 (b) features fixed residues, including *I* at position 7, with strong recommendations for *W* at position 2, *I* at position 4, and *R* at positions 8 and 11. Position 1 and 10 allow *R* to *W* substitutions, whereas position 9 permits *V* to be replaced with *Y*, *W*, or *T*.

For Motif 3 (”KIWKKJLKKIIKPVK”) shown in Figure 13 (c), *K* is strongly preferred at positions 1, 4, 5, 8, 9, 12, 13, and 15. Exceptional conservation occurs at positions 4, 5, 10, and 15, whereas other positions allow substitutions with lower-frequency residues.

#### 3.5.3 Wet Lab Experiments

We conducted wet-lab experiments of MIC determination and hemolytic activity assay on human RBCs. The MIC determination against *Escherichia coli* ATCC 25922, *Staphylococcus au-reus* ATCC 25923, and methicillin-resistant *S. aureus* (MRSA, ATCC 43300) in accordance with the Clinical and Laboratory Standards Institute (CLSI) broth microdilution method M07–A11 [61]. The hemolytic activity of the peptides was assessed against human red blood cells (RBCs) according to a protocol based on Sabo et al. [62].

As shown in Table 7, we removed the longest sequence, Seq7, from Table 6, retaining 11 sequences for wet lab experiments against EC, SA, and methicillin-resistant *Staphylococcus aureus* (MRSA). Furthermore, we examined the 10% hemolytic concentration (HC_10_) using human red blood cells. In contrast to HC_50_, which serves as a benchmark for toxicity potency, HC_10_ was selected as a more stringent safety threshold. As shown in Table 7, no hemolysis was observed at concentrations up to 256 *µg/mL*, the highest concentration tested. Therefore, the therapeutic index (TI), calculated as the ratio of the hemolytic concentration to the MIC, is estimated to be at least 16, indicating a favourable safety margin. Hence, on the basis of the results in Table 7, it can be concluded that, with the exception of Seq2, Seq3 and Seq10, all the other 8 sequences have potential as novel drug candidates.

**Table 7:** Antimicrobial activity and toxicity scores of selected sequences subjected to wet laboratory experiments.

| ID | Sequence | EC<sub>MIC</sub> | SA<sub>MIC</sub> | MRSA<sub>MIC</sub> | Toxic dose | TI<sub>EC</sub> | TI<sub>SA</sub> | TI<sub>MRSA</sub> |
| --- | --- | --- | --- | --- | --- | --- | --- | --- |
| Seq1 | RRWWRWWRI-NH2 | 16 | 16 | 16 | 256 | 16 | 16 | 16 |
| Seq2 | YRWWRWWWR-NH2 | 16 | 8 | 16 | 64 | 4 | 8 | 4 |
| Seq3 | WRTRRRWRY-NH2 | 64 | 64 | 128 | >256 | 4 | 4 | 2 |
| Seq4 | KRRWKTWWW-NH2 | 32 | 16 | 32 | >256 | 8 | 16 | 8 |
| Seq5 | RWKWRWRSW-NH2 | 8 | 16 | 32 | >256 | 32 | 16 | 8 |
| Seq6 | KWRRFKWWL-NH2 | 16 | 16 | 32 | >256 | 16 | 16 | 8 |
| Seq8 | RWRRWKYWY-NH2 | 16 | 32 | 64 | >256 | 16 | 8 | 4 |
| Seq9 | KRRYRWIWI-NH2 | 16 | 16 | 32 | 256 | 16 | 16 | 8 |
| Seq10 | FRWRIVVIRYRR-NH2 | 16 | 32 | 16 | 64 | 4 | 2 | 4 |
| Seq11 | LWRRWYRWI-NH2 | 16 | 16 | 32 | 256 | 16 | 16 | 8 |
| Seq12 | YKKRWRLWI-NH2 | 16 | 32 | 64 | >256 | 16 | 8 | 4 |

All MIC values for EC, SA, and MRSA and Toxic dose are reported in $\mu\text{g/mL}$ .
Toxic dose is defined as the concentration at which hemolysis of human red blood cells exceeds 5%.

## 4 Conclusion

We propose a classifier guidance diffusion model called ClsDiff-AMP30 for progressively inferring higher AMP score sequences with lengths smaller than 30 from sampled Gaussian noise and achieve a generation accuracy of 66%. ClsDiff-AMP30 consists of a noise predictor that can denoise a given noisy token progressively to a clean token and a noisy classifier that aims to refine a given noisy token to possess a higher AMP score and finally reverse the classifier-refined clean token to an AMP sequence. To prove the efficiency of the ClsDiff-AMP30 model, we generated 10,000 sequences each time and developed an RFClassifier-AMP30 model to classify the generated sequences as AMP or not. To find the best performance model for the proposed ClsDiff-AMP30 model and the self-developed RFClassifier-AMP30 model, we performed a series of hyperparameter searches with the grid search method. Furthermore, we conducted a series of experiments to prove the generation power of the ClsDiff-AMP30 model on the basis of generation AMP sequences (GeneAMP), real AMP sequences (RealAMP), random sequences (RndSeq), and random distinct peptides (RndPep). With the four datasets, we conducted a series of qualitative experiments on the basis of length distribution, amino acid composition, feature combinations, and physicochemical characteristics. To explore the robustness of the generative power of ClsDiff-AMP30, we also conducted experiments on the ClsDiff-AMP30 model based on different data sizes, changing the input Gaussian noises to gamma and uniform noises, and performed an ablation study to explore the generation efficiency of the ClsDiff-AMP30 model with and without the classifier guidance module. We subsequently screened 12- and 130-generation AMP sequences to visualize the 3D structures and find their motifs, respectively. Finally, wet laboratory evaluation of the 11 selected sequences, out of an initial 12 with the longest sequence excluded for cost reasons, revealed that all the sequences displayed high antimicrobial activity and low hemolytic activity against the three tested bacterial strains, confirming that they are promising candidates.

## 5 Author contributions statement

S.W.I.S. and F.X.C.V. conceived the study. J.Y. and J.C. prepared the datasets and conducted the experiments. J.Y. designed the methods, analysed the results, and drafted the manuscript. Y.L. implemented the data analysis of the results. I.F.L. and Z.L. completed the wet-lab experiments.

M.Z. and X.W. revised the writing. S.W.I.S., F.X.C.V., and X.Z. supervised the work. S.W.I.S. and F.X.C.V. finalized the manuscript. All the authors read and approved the final manuscript. F.X.C.V., S.W.I.S., and J.Y. acquired funding. The authors declare that they have no conflicts of interest.

## Acknowledgements

The work is supported by the Government of Canada’s New Frontiers in Research Fund (NFRF) (NFRFE-2021–00913), the Postdoctoral Fellowship Program of CPSF under Grant GZC20233322, and the Postdoctoral Talent Special Program. J.Y. was supported by the NFRF, the CPSF, and the Postdoctoral Talent Special Program. J.C. was the recipient of the Macau Polytechnic University graduate scholarship. W.X. was supported by the General Program of the Natural Science Foundation of Chongqing under Grant CSTB2024NSCQ-MSX0479, the Chongqing Postdoctoral Foundation Special Support Program under Grant 2023CQBSHTB3119, and the China Postdoctoral Science Foundation under Grant 2024MD754244. The funders had no role in the study design, data collection, interpretation, or decision to submit the work for publication.

## Data availability

The peptide data used in this study come from DBAASP [34] (https://dbaasp.org/home). The data and scripts for data preparation to reproduce the experiments are available at https://github.com/jieluyan/ClsDiff-AMP30. (They will be made accessible upon manuscript acceptance.) The final prediction models trained in this study are also available in the previous link. The web server used to access this prediction method directly is https://app.cbbio.online/ampep/home.

